# Differences in Conformational Sampling and Intrinsic Electric Fields Drive Ion Binding in Telomeric and TERRA G-Quadruplexes

**DOI:** 10.1101/2023.08.10.552810

**Authors:** Marcelo D. Polêto, Justin A. Lemkul

## Abstract

The formation of G-quadruplexes (GQs) occurs in guanine-rich sequences of DNA and RNA, producing highly stable and structurally diverse noncanonical nucleic acid structures. GQs play crucial roles in regulating transcription, translation, and replication; and maintaining the genome, among others, thus changes to their structures can lead to diseases such as cancer. Previous studies using polarizable molecular dynamics simulations have shown differences in ion binding properties between telomeric and TERRA GQs despite architectural similarities. Here, we used volume-based metady-namics and repulsive potential simulations in conjunction with polarizable force fields to quantify the impact of ion binding on GQ dynamics and ion binding free energies. Furthermore, we describe how GQs exert electric fields on their surroundings to link dynamics with variations in electronic structure. Our findings provide new insights into the energetic, physical, and conformational properties of GQs and expose subtle, but important, differences between DNA and RNA GQs with the same fold.

## Introduction

G-quadruplexes (GQs) are noncanonical nucleic acid structures that form in DNA and RNA sequences that are rich in guanine. GQs are composed of a core structure of Hoogsteen hydrogen-bonded guanine tetrads and loop structures that vary in length and sequence. The tetrad bases are oriented such that their carbonyl oxygen (O6) atoms are directed inward, creating an electronegative channel (O6-core) that is stabilized by monovalent cations. GQ sequence motifs are prevalent in various genomic regions such as promoters,^1,2^ origins of replication,^3,4^ telomeres,^5^ and untranslated regions in mRNA.^6,7^ The presence of GQs in these regions suggests a crucial role in modulating gene expression and preserving genomic integrity. Hence, GQs are considered as potential targets for drug design.

Telomeric DNA is a guanine-rich region with d(TTAGGG) repeats. As such, telomeric DNA can form both unimolecular^8^ and bimolecular^9^ GQ structures that are the substrate for telomerase activity.^5^ Stabilization of telomeric GQs by small molecules has been shown to inhibit telomerase activity, leading to cell death.^10,11^ The transcription of the complementary telomeric strands produces telomeric repeat-containing RNA (TERRA) sequences that regulate heterochromatin formation, telomerase activity, and homologous recombination.^12^ Previous work has shown that TERRA can form GQ structures^13^ with the same bimolecular, parallel architecture as telomeric GQs.^9^ In addition, TERRA GQs are recognized by telomeric repeat-binding factors and the disrupting these complexes leads to increased telomeric aberrations and decreased telomeric length.^14^ Considering that telomerase is overactive in several cancer types and telomeric integrity is linked to cell death, telomeric and TERRA GQs are considered potential drug targets.^7,12,14,15^

Molecular dynamics (MD) simulations have proven to be a useful tool for investigating the conformational dynamics of GQs. A key challenge in accurately modeling GQs is accounting for ion-ion and ion-GQ interactions, which has limited the success of GQ simulations performed with nonpolarizable (additive) force fields.^16–18^ Gkionis et al. ^19^ demonstrated that neglecting explicit electronic polarization terms can result in an overestimation of ion-ion repulsion by additive force fields. In contrast, our group^20–25^ and others^17^ have used the Drude polarizable force field to study the conformational dynamics of a variety of DNA and RNA GQs, observing core K^+^ ion retention, unbiased K^+^ binding/unbinding events, distinct coordination and dehydration properties of monovalent ions around GQs, and distinct ion distributions around DNA and RNA GQs. In one such example, we found that, despite sharing the same fold and analogous sequences, telomeric and TERRA GQs exhibit different dynamic behaviors.^25^

Here, we build on the previous work by employing enhanced sampling techniques to investigate the tetrad core dynamics of both telomeric and TERRA GQs, with the goal of describing how these dynamics influence ion binding. A key question that remains unanswered is whether intrinic fluctuations in GQ structure help recruit ions, or whether ions bind and subsequently perturb the structure of GQs in any way. Beyond this question, the binding free energies of ions to GQs remain unknown, we hypothesize that this property is linked to induced polarization between the GQ and ions, and among the ions themselves. We applied cylinder-based metadynamics simulations to calculate ion binding free energies. Lastly, we explored the organization of electric fields surrounding these GQs and how they impact ion binding. Our approach provides new insights into the specific contribution of electronic and conformational properties in GQs that can be used for future studies into protein-GQ interactions and drug design efforts.

## Results and discussion

### Overview of GQ structure and nomenclature

Lemkul ^25^ previously simulated telomeric and TERRA GQ via unbiased simulations using the Drude-2017 polarizable force field.^26–28^ Here, we used the equilibrated coordinates of each replicate of the telomeric and TERRA GQ systems in that study as starting coordinates for our own biased simulations described below. The unbiased trajectories reported by Lemkul ^25^ were also analyzed in this work and served as reference for our new simulations.

Some important naming conventions are illustrated in Figure 1. Tetrads 1, 2 and 3 are named T1, T2 and T3, respectively. Being bimolecular GQs, T1 comprises residues 3 and 9 from each GQ segment. Similarly, T2 contains residues 4 and 10, and T3 contains residues 5 and 11. The K^+^ ions coordinated between T1 and T2 is defined as K1, while K2 is coordinated between T2 and T3. Moreover, unbiased simulations reported by Lemkul ^25^ described bulk K^+^ binding events close to T1 and T3. Those binding sites are defined here as S1 and S3, respectively.

**Figure 1:**
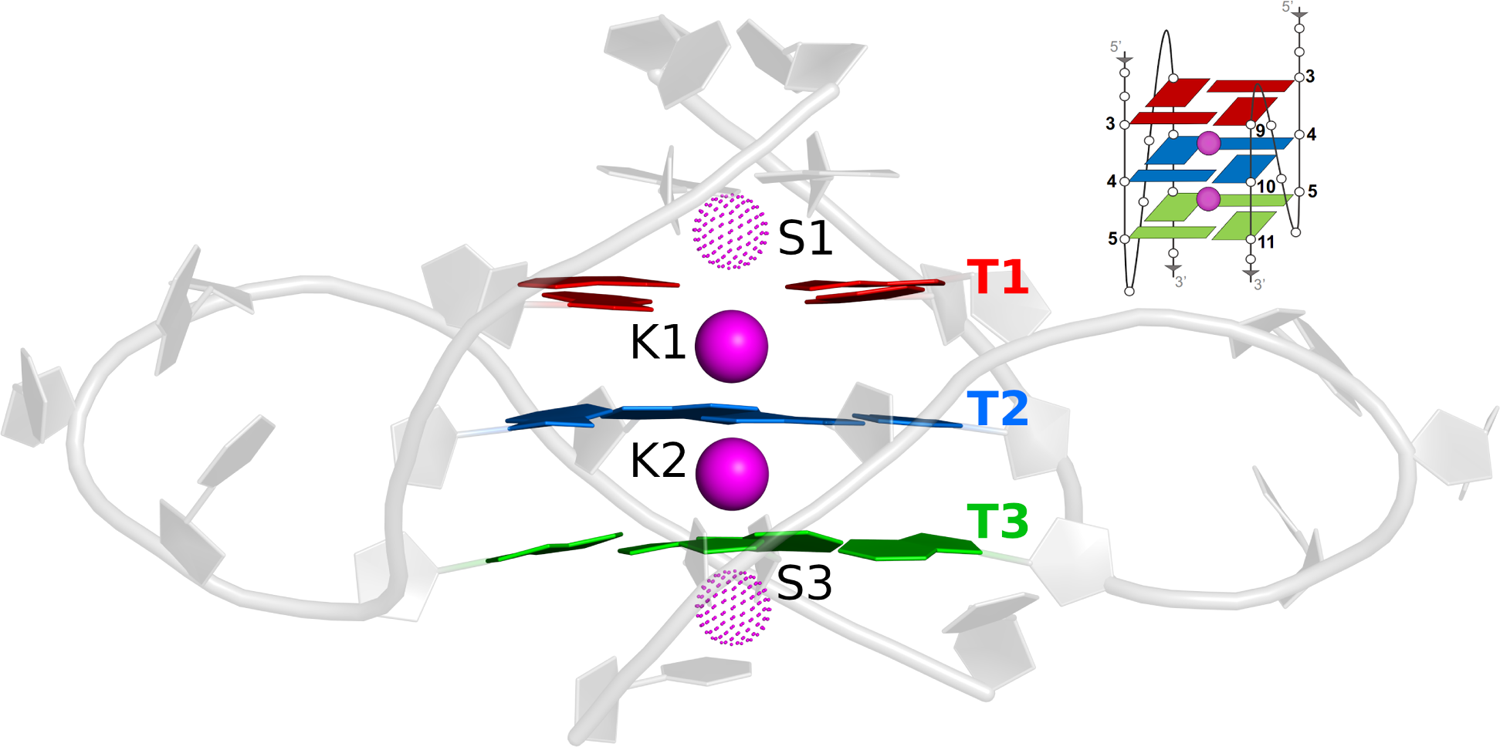
Structural definitions used for telomeric and TERRA GQs in this work. Tetrads 1, 2 and 3 (T1, T2, and T3) are shown in red, blue and green, respectively. K1 and K2 are the coordinated K^+^ ions in the folded GQ structure. S1 and S3 are binding sites for bulk K^+^ ions.

### GQ core dynamics

The unbiased simulations of TERRA and telomeric GQs reported by Lemkul ^25^ detailed reversible binding events of K^+^ at binding site S3. A meta-analysis of those simulations revealed an intrinsic difference in the tilting direction of the tetrad guanine bases in both of these GQs. Moreover, K^+^ binding at S3 seemed to modulate the base tilting motion, but a rigorous conclusion of whether these tilting motions were caused by ion binding could not be drawn from the unbiased simulations alone. To investigate those differences, we performed a new set of simulations of both GQs with a repulsive potential between K1/K2 and all bulk K^+^ ions to prevent ion binding at S1 and S3. Doing so allowed us to analyze the intrinsic dynamics of the bases without influence of bulk K^+^ ions. In contrast, we selected the segments of the unbiased trajectories across replicas in which K^+^ were bound at S3 to generate an ensemble of the bound states of each system, thus allowing a direct comparison between bound and unbound states.

We calculated the average tilting angle of guanines in each tetrad relative to its tetrad plane (Figure 2A and 2B). For the telomeric GQ, the tilting angles were mostly positive (tilted towards S1) in the unbound state, with averages of 23 *±* 10*^◦^*, 29 *±* 9*^◦^*, and 32 *±* 10*^◦^* for T1, T2 and T3, respectively. For the TERRA GQ, the tilting angles were mostly negative (tilted towards S3) in the unbound state, but the magnitude of tilting was lower than the telomeric GQ tetrads, with average tilting angles for T1, T2 and T3 of −11 *±* 10*^◦^*, −23 *±* 9*^◦^*, and −8 *±* 10*^◦^*, respectively.

**Figure 2:**
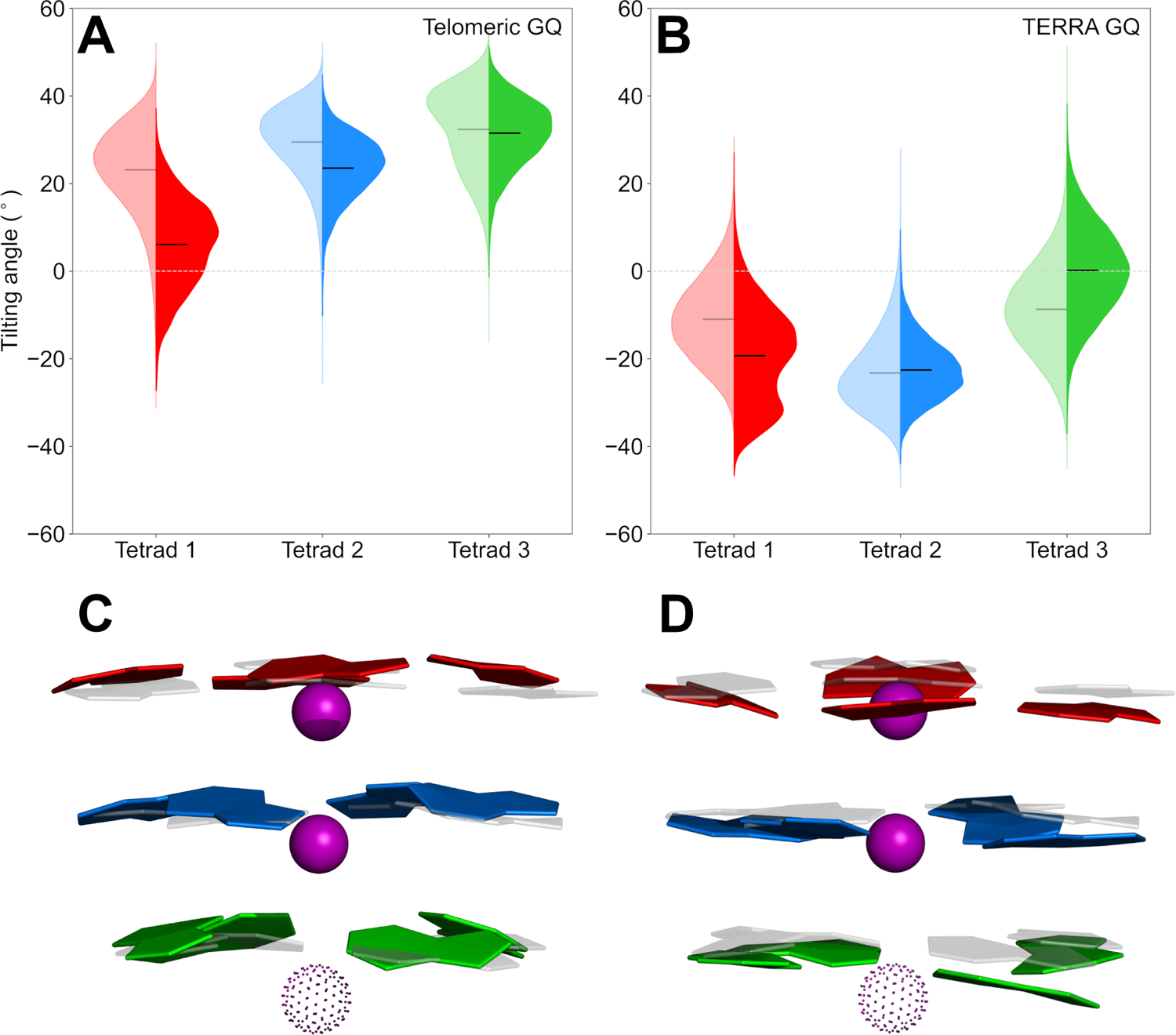
(A-B) Violin plots of the average tilting angles for base tetrads of telomeric and TERRA GQs. Bound states are shown as opaque right-sided distributions, while the unbound states are shown as translucent left-sided distributions. Mean values are shown in black and grey lines inside the distributions. The dotted line at 0*^◦^*marks the C1’ plane of each tetrad. (C-D) Superposition of X-ray structure (transparent) and a representative structure of telomeric and TERRA GQs with tilted tetrad bases. The S3 binding site is shown as a dotted sphere.

Considering the X-ray structures of the telomeric and TERRA GQs, T1 and T2 are almost planar in both systems, with base tilting angles smaller than 5*^◦^*, while T3 is tilted positively (towards S1) about 25*^◦^*. Nevertheless, the conformational ensemble from our simulations suggests that the tetrad cores of both telomeric and TERRA GQs are not only tilted, but tilt in opposite directions prior to ion binding events. In fact, the NMR structural ensemble of TERRA obtained in K^+^ solution (PDB 2KBP) by Martadinata and Phan ^29^ (average tilting angles of −11 *±* 4*^◦^*, −17 *±* 3*^◦^*, and −20 *±* 3*^◦^* for T1, T2 and T3, respectively) qualitatively agrees with the base tilting direction observed in our simulations.

Greater perturbations to tilting dynamics in T3 bases might be expected upon ion binding at S3, but this phenomenon was not observed for the telomeric GQ. T3 was mostly unaffected (31 *±* 7*^◦^*), while T1 and T2 had a more pronounced decrease in their average tilting angle (6 *±* 10*^◦^*and 23 *±* 7*^◦^*, respectively). For TERRA GQ, T3 was the most affected tetrad and its tilting angle averaged around 0 *±* 10*^◦^* in the bound state. Interestingly, T1 dynamics were affected by K^+^ binding in both systems even though it is the furthest tetrad from S3. However, while T1 tilting angle decreased in the telomeric GQ, in the TERRA GQ, T1 increased its tilting angle.

Based on the different tilting motions of the telomeric and TERRA GQs, we investigated the ribose puckering of each tetrad nucleotide by calculating the pseudorotation angle defined by Altona and Sundaralingam ^30^. In this convention, angles between 0-36*^◦^* and 144-180*^◦^* correspond to C3’-endo and C2’-endo conformations, respectively. Figure 3 shows per-residue sugar puckering for both GQ systems in their bound and unbound states.

**Figure 3:**
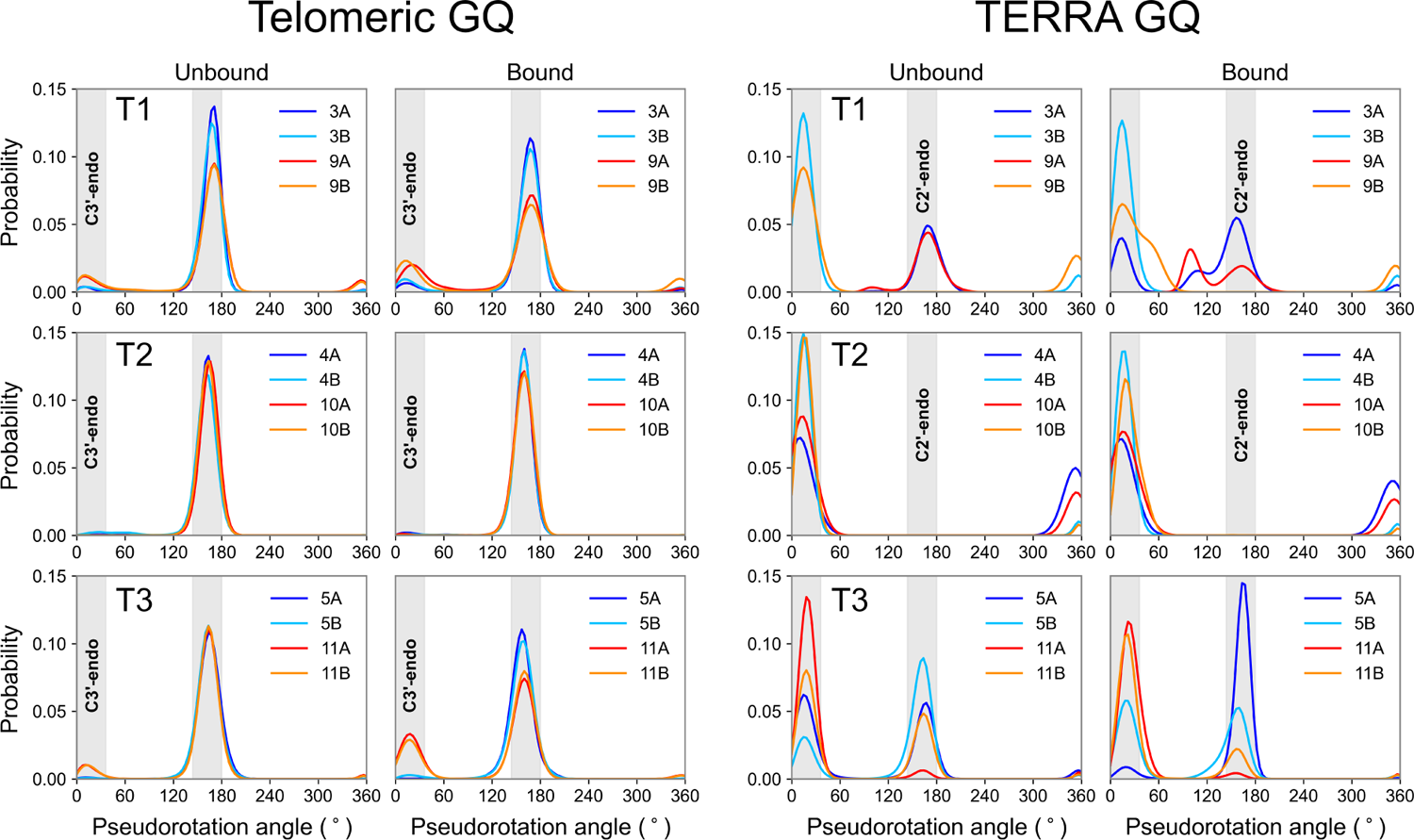
(Deoxy)ribose puckering distributions for telomeric and TERRA GQs in their bound and unbound states. Regions corresponding to C2’-endo and C3’-endo conformations are highlighted in gray. Tetrad residues for chains A and B are shown in cold (blue and cyan) and warm (red and orange) colors, respectively.

Guanine residues in all three tetrads of the telomeric GQ predominantly sampled C2’-endo conformations in the unbound state, with relative abundances of 73%, 87%, and 79% for T1, T2 and T3, respectively. K^+^ binding did not seem to impact this trend, but it slightly increased minor C3’-endo populations in T1 and T3 in the bound state. The preference for C2’-endo ribose pucker is in line with the expected deoxyribose behavior in DNA.

The TERRA GQ, however, manifested more heterogeneous sampling among the tetrads, in line with previous reports for RNA GQs.^31,32^ An asymmetric behavior among the tetrad bases in T1 and T3 was also observed. In the unbound state, T1 G3 and G9 of chain A (G3A and G9A, respectively) sampled C3’-endo conformations, whereas G3 and G9 of chain B (thus G3B and G9B) sampled C2’-endo conformations. In T3, G11A favored C3’-endo conformations, but the rest of its tetrad guanines sampled both C3’- and C2’-endo conformations. T2 sugars were consistently in C3’-endo conformations. Upon K^+^ binding to S3, guanine residues in T2 maintained the overall puckering pattern, but guanines in T1 sampled intermediate states between C2’-endo and C3’-endo, and the T3 G5A distribution shifted to C2’-endo and the G11B distribution shifted to C3’-endo, maintaining the overall puckering abundances at the same levels. It is noteworthy that the presence of C2’-endo conformations in TERRA GQ tetrads observed here were also reported by Martadinata and Phan ^29^, suggesting the structural ensemble captured by our simulations is supported by experimental evidence. Together, our results suggest that even though telomeric and TERRA GQs have the same fold, their core structural properties are fundamentally different, including their sensitivity to the effects of K^+^ binding.

### Polarization response upon binding

Given the correlation between K^+^ binding events and structural perturbations in the GQ cores, we investigated the electronic polarization response of the GQs upon ion binding. To do so, we calculated the dipole moment of guanine bases in each tetrad for both bound and unbound states (Table 1). The use of a repulsive potential (see Methods) enabled us to prevent ion binding at the S1 and S3 sites, thus more comprehensively sampling unbound states that are otherwise rare in unbiased simulations.^25^ Overall, our present results follow the trend reported previously, but some noticeable differences must be highlighted.

**Table 1:** Total dipole moments (in Debye) of tetrad guanines in telomeric and TERRA GQ in both bound and unbound states.

| **Dipole Moment (D)** |  |  |
| --- | --- | --- |
| **Telomeric GQ** | **Unbound** | **Bound** |
| **T1** | $8.8 \pm 0.5$ | $8.8 \pm 0.5$ |
| **T2** | $9.3 \pm 0.4$ | $9.3 \pm 0.4$ |
| **T3** | $9.0 \pm 0.5$ | $9.3 \pm 0.5$ |
| **TERRA GQ** | **Unbound** | **Bound** |
| **T1** | $8.7 \pm 0.5$ | $8.8 \pm 0.5$ |
| **T2** | $9.1 \pm 0.4$ | $9.1 \pm 0.4$ |
| **T3** | $9.1 \pm 0.5$ | $9.2 \pm 0.5$ |

The previous unbiased simulations sampled both bound and unbound states, and the guanine base dipole moments reported there reflect an average of those two states. Here, by preventing ion binding to S3, we were able to clearly distinguish the dipole properties of these GQs with and without auxiliary ions bound at S1 and S3. In the unbound state, the average guanine base dipole moments of T1, T2 and T3 in the telomeric GQ were 8.8 *±* 0.5 D, 9.3 *±* 0.4 D and 9.0 *±* 0.5 D, respectively. For the TERRA GQ, the average base dipole moments were 8.7 *±* 0.5 D, 9.1 *±* 0.4 D and 9.1 *±* 0.5 D for T1, T2 and T3, respectively. Upon ion binding to S3, the average guanine base dipole moment of T3 increased to 9.3 *±* 0.5 D and 9.2 *±* 0.5 D for telomeric and TERRA GQs, respectively, which is expected due to the proximity of the tetrad to the ion binding site. It is interesting to note that T2 was slightly more polarized in telomeric GQ relative to the TERRA GQ. Thus, we directly quantified the differences in how the electronic properties of these GQs respond to perturbations due to K^+^ binding.

Because these polarization differences likely impact the nonbonded interaction energy (an approximation of the enthalpy of binding) between the GQs and K^+^ binding at S3, we calculated the role of explicit polarization in our simulations (Table 2). First, we calculated interaction energies between the tetrad bases and the K^+^ ion binding at S3, defining a “bound” state as having a K^+^ ion within 5 Å of K2 in the tetrad core. We subsequently converted the Drude coordinates and topologies to their CHARMM36 equivalents by deleting the Drude particles and lone pairs. Doing so allowed us to compute interaction energies with the CHARMM36 force field using equivalent configurations. The use of an additive force field here models the absence of explicit polarization response while still using a well-calibrated force field that shares similar parametrization theory and methodology. Thus, by using the same trajectory for both cases, this approach allow us to directly compare interaction energies from each force field.

**Table 2:** Average interaction energies (in kcal/mol) between K^+^ ions binding at S3 and GQ tetrad bases.

|  | All tetrads |  | Tetrad 3 only |  |
| --- | --- | --- | --- | --- |
|  | Drude-2017 FF | CHARMM36 FF | Drude-2017 FF | CHARMM36 FF |
| Telomeric GQ | $-26 \pm 5$ | $-15 \pm 4$ | $-25 \pm 5$ | $-15 \pm 4$ |
| TERRA GQ | $-30 \pm 4$ | $-21 \pm 3$ | $-29 \pm 4$ | $-20 \pm 3$ |

Considering the contribution of all tetrad bases, the interaction energies calculated using the additive topologies for telomeric and TERRA GQs were −15 *±* 4 and −21 *±* 3 kcal/mol, respectively. By considering explicit polarization in the Drude model, the interaction energies were stronger, −26 *±* 5 and −30 *±* 4 kcal/mol for telomeric and TERRA GQs, respectively. This *≈*10 kcal/mol decrease in both systems suggests a considerable enthalpic contribution arising from the multibody polarization effects upon K^+^ binding. Interestingly, the same *≈*10 kcal/mol difference was observed when we considered only the contribution of T3 to the interaction energies, suggesting that electronic polarization response upon K^+^ binding to the GQs is strongly localized to T3. We observed a similar phenomenon with respect to multibody effects among core K^+^ ions and tetrad guanine bases in the *c-kit1* GQ.^20^ In that study, we found that polarization response upon auxiliary ion binding increased the strength of core K^+^ interaction energy closest to the binding ion, with a sharp decay in this effect for the other tetrads. Thus, these properties appear to be common among different GQs.

It is important to highlight that K^+^ interacted systematically more strongly with the TERRA GQ than the telomeric GQ. By quantifying the interaction energies considering and neglecting explicit electronic polarization, we conclude that the base tilting manifested by the GQ tetrad core plays an important role in the interaction strength with binding ions. That is, the inward tilting of tetrad 3 bases upon K^+^ binding appears to enable stronger binding than in the case of the telomeric GQ. We discuss the nature of these differences in greater detail below.

### K^+^ binding free energy

We previously reported ion binding events to the telomeric and TERRA GQs that occurred over hundreds of nanoseconds,^25^ making it difficult to accurately assess ion binding affinity due to infrequent exchanges. To overcome this issue and quantitatively assess ion affinity, and to further probe the extent to which the conformational dynamics of TERRA and telomeric GQs impact ion binding free energy, we performed well-tempered cylinder metadynamics simulationsusing the ion position inside the cylinder as a collective variable (see Methods). The resulting two-dimensional (2D) free energy surfaces (FES) are shown in Figure 4. For the telomeric GQ, there was a clear global free energy minimum at binding site S3 (*x*=0, *y*=0 and *z*=-4.50 Å), along with two local minima close to the binding site (Figure 4A,C). For the TERRA GQ, a global free energy minimum was also observed at S3 (*x*=0, *y*=0 and *z*=-6.50 Å), with no other clear minima (Figure 4B,D).

**Figure 4:**
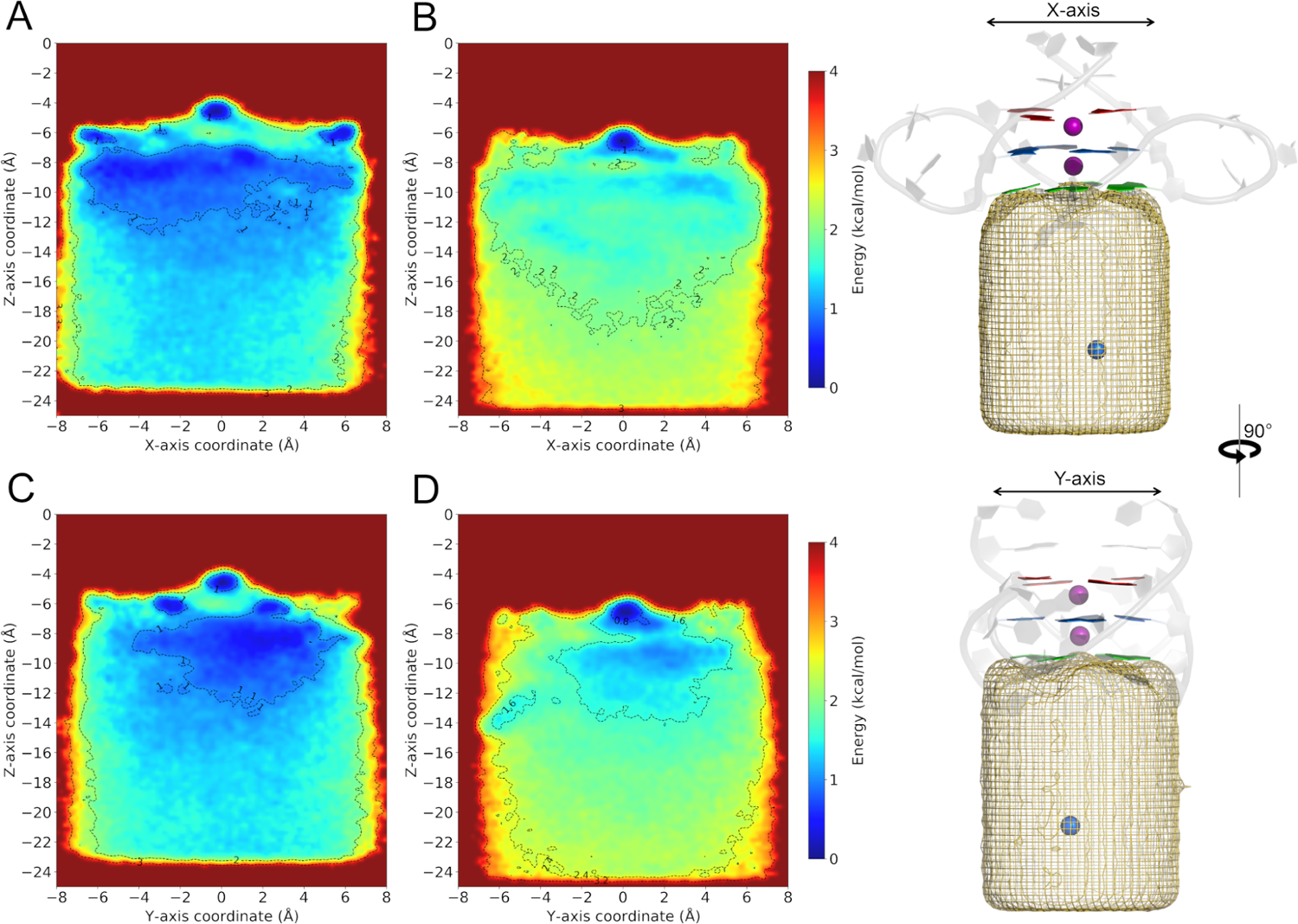
2D free energy surfaces associated with K^+^ binding. The *x*-axis projections for telomeric and TERRA GQ are shown in panels A and B, respectively, while *y*-axis projections are shown in panels C and D. The definitions of the *x*- and *y*-axis projections are also shown in the context of the telomeric GQ structure, with the cylindrical volume illustrated as a mesh.

We assume that K^+^ ions beyond 15 Å from the global minima (along the *z*-axis) behave like bulk ions. The free energy values in the bulk region for TERRA GQ are higher than their counterparts for the telomeric GQ, suggesting a more favorable ion binding free energy (which we denote as Δ*G_MET AD_*) to the TERRA GQ. However, these projections of free energy onto a 2D surface might not represent the real 3D binding pathways, from which we could more confidently quantify Δ*G_MET AD_*.

To resolve these pathways, we employed MEPSAnd,^33^ a minimum energy path surface analysis to detect possible binding pathways from the bulk region to the global minimum in conjunction with a 3D FES based on the *x*-, *y*- and *z*-axis coordinates of the cylinder volume. First, MEPSAnd detects local minima in the FES space and later uses these minima as reference to trace pathways connecting 2 specific minima. The rationale is that binding events tend to follow minimum energy pathways.

For each system, we selected 10 pathways connecting local minima in the bulk region to the global minimum, as shown in Figure 5. To estimate the ion binding free energies (Δ*G_MET AD_*) of each system, we averaged the free energy values of the bulk region from the selected paths. For telomeric GQ, Δ*G_MET AD_* was −2.6 *±* 0.2 kcal/mol, while for the TERRA GQ, Δ*G_MET AD_* was −3.4 *±* 0.2 kcal/mol. By calculating the correction to the binding free energy due to the change in the translational entropy^34,35^ of the binding K^+^ in the bulk region, we obtain a standard free energy of binding (Δ*G*^0^) of −2.4 *±* 0.2 kcal/mol and −3.1 *±* 0.2 kcal/mol for telomeric and TERRA GQs, respectively.

**Figure 5:**
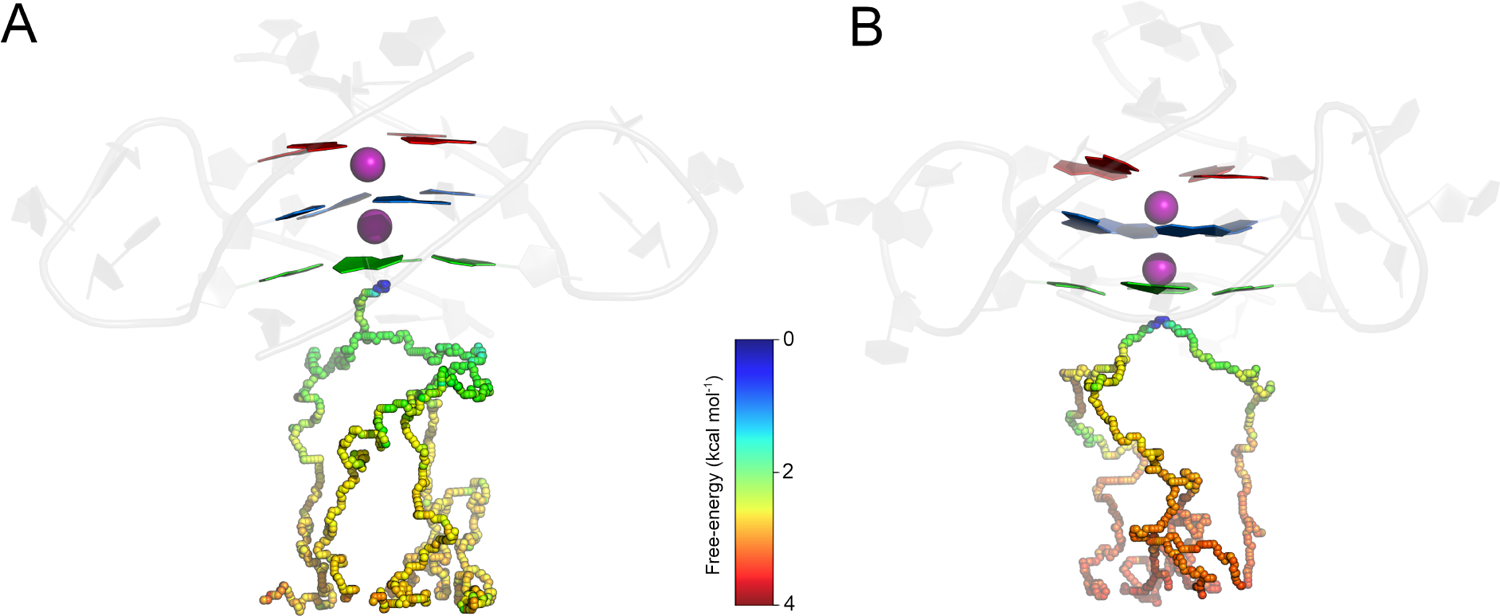
K^+^ binding pathways for (A) telomeric and (B) TERRA GQs colored as a function of free energy values obtained from the 3D FES.

Overall, our results are in line with previous qualitative observations of more persistent K^+^ binding at S3 for the TERRA GQ,^25^ suggesting that K^+^ ions more favorably to bind to S3 in the TERRA GQ than the telomeric GQ. Our present results suggest a Δ*G*^0^ that is stronger by 0.7 kcal/mol, which is in line with the stronger interaction energies reported above for TERRA GQ. Hence, these results suggest that the tetrad core dynamics and an electronic polarization response act together to recruit K^+^ ions to S3.

### Electric fields exerted by GQs

Intrigued by the well-defined global free energy minimum at the S3 binding site in both systems, we investigated the electrostatic forces stabilizing the binding of K^+^ ions. To this end, we employed TUPÃ^36^ to calculate the electric fields (*E^→^*) exerted by the GQs across their O6-core axis for unbound states (snapshots with no bound K^+^ bound at S1 or S3, Figure 6). To facilitate our analysis, we aligned the O6-core axis to the Cartesian *z*-axis, and the plane formed by C1’ atoms of T2 was used to define *z*=0. As explained in the Methods, we selected residues 3 to 11 to act as the environment set.

**Figure 6:**
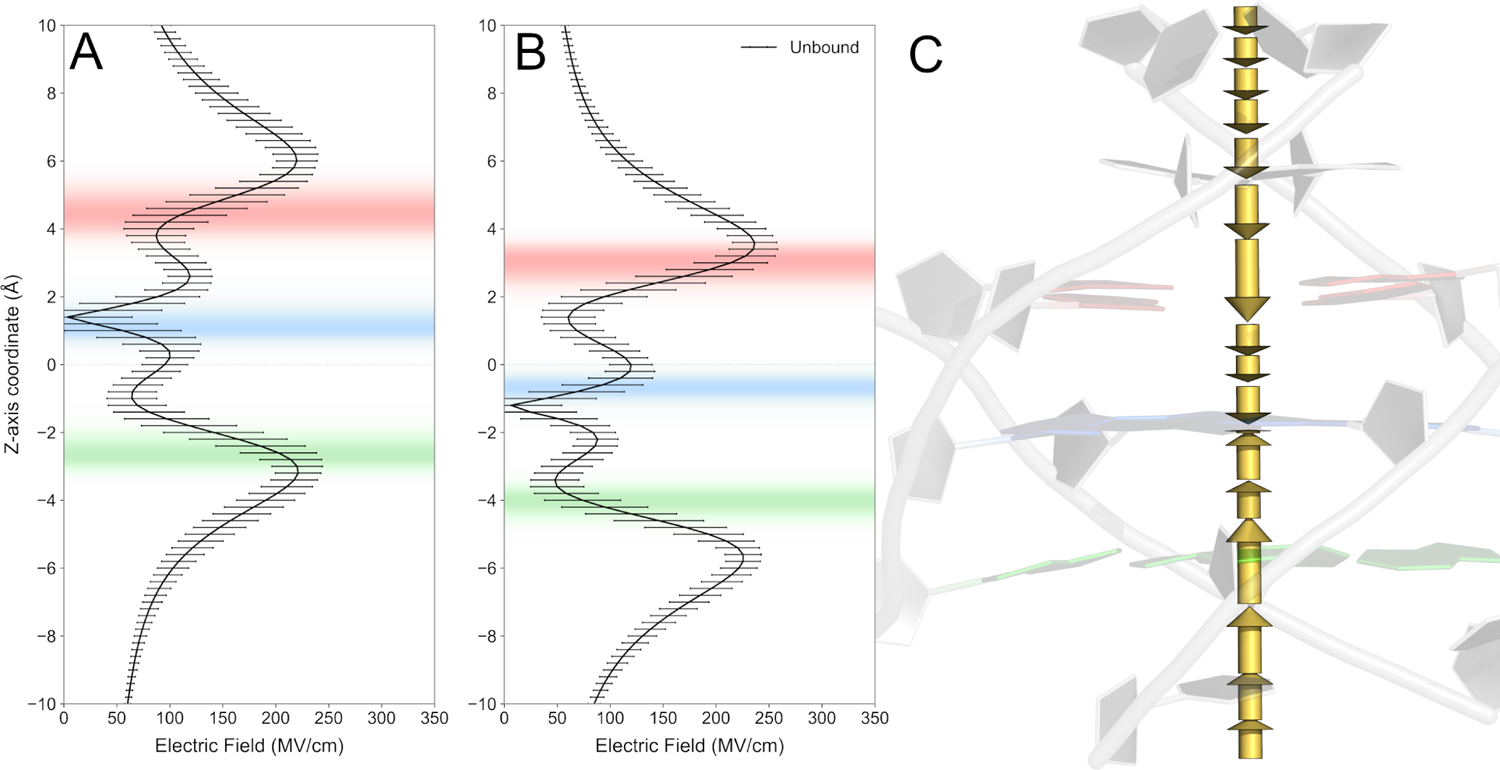
Values of *|E^→^_Z_|* along the O6-core axis for the (A) telomeric and (B) TERRA GQs for unbound states. The *z*-axis locations of T1, T2 and T3 guanine O6 atoms in are shown as distributions in red, blue and green, respectively. In (C), the direction of field vectors across O6-core axis is illustrated with arrows with lengths proportional to the magnitudes of *E^→^_Z_*at those points in space.

Across biased and unbiased simulations of both systems, the greatest contribution to *E^→^*came from the axis orthogonal to the tetrad plane (*E^→^_Z_*). This outcome is expected due to the symmetry axis of GQ tetrads, which, on average, results in a cancellation of field contributions in the tetrad plane (*E^→^_X_*and *E^→^_Y_*), yielding a resultant orthogonal to the tetrad plane. Figures S2 and S3 show *E^→^_X_*and *E^→^_Y_*across the O6-core axis.

In addition, the stacking of three tetrads creates a symmetry plane at the middle tetrad, canceling the orthogonal contributions close to T2. For the telomeric GQ, *E^→^_Z_*is very close to zero precisely at T2, whereas the minimum *E^→^_Z_*value for the TERRA GQ is slightly shifted towards T3. This difference is likely related to the different base tilting profiles observed for the two GQs. Overall, *E^→^_Z_*is also weaker between tetrads, where K1 and K2 are coordinated, with a slightly stronger field between T1 and T2.

It is also interesting to note that the maximum values of *E^→^_Z_*occur close to ion binding sites S1 and S3, with some relevant differences between telomeric and TERRA GQs. Considering the tetrad motions upon binding, we can generalize that S1 and S3 are located *≈*2Å from T1 and T3, respectively. Previous work using the Drude force field to study GQs has described persistent binding of bulk ions to binding sites close to tetrads and aligned to the O6-core axis in unbiased MD simulations,^21–24^ with no clear preference for which side of the GQ such binding occurred. Here, we show how the tetrad core architecture of GQs organizes strong electric fields at those binding sites, driving ion binding.

Interestingly, the field profiles along the O6-core axis are very similar between the telomeric and TERRA GQs, but shifted in opposite directions. At S3, *E^→^_Z_*is *≈*160 MV/cm for the telomeric GQ in unbound states, whereas *E^→^_Z_*is *≈*200 MV/cm for the TERRA GQ. The opposite is true for S1: the telomeric GQ exerts a stronger field (*≈*200 MV/cm) than the TERRA GQ (*≈*160 MV/cm). Together, these results suggest that tetrad dynamics shift the field profiles differently along the O6-core axis in the unbound state, allowing a stronger field at S3 for TERRA GQ and a stronger field at S1 for telomeric GQ.

Considering the tetrad tilting motions upon K^+^ binding at S3, we calculated *E^→^_Z_*along the O6-core axis for snapshots with bound K^+^ ions using the same conventions used for the unbound states. As shown in Figure 7, ion binding modulates the field profiles differently. Whereas *E^→^_Z_*values between T2 and T3 decreased upon ion binding for both system, there was a slight increase of *E^→^_Z_*between T1 and T2 for the TERRA GQ. Moreover, both GQs exerted a stronger *E^→^_Z_*at binding site S3 (*≈*230 MV/cm). Such a strong field explains the existence of a clear free energy global minimum at this location.

**Figure 7:**
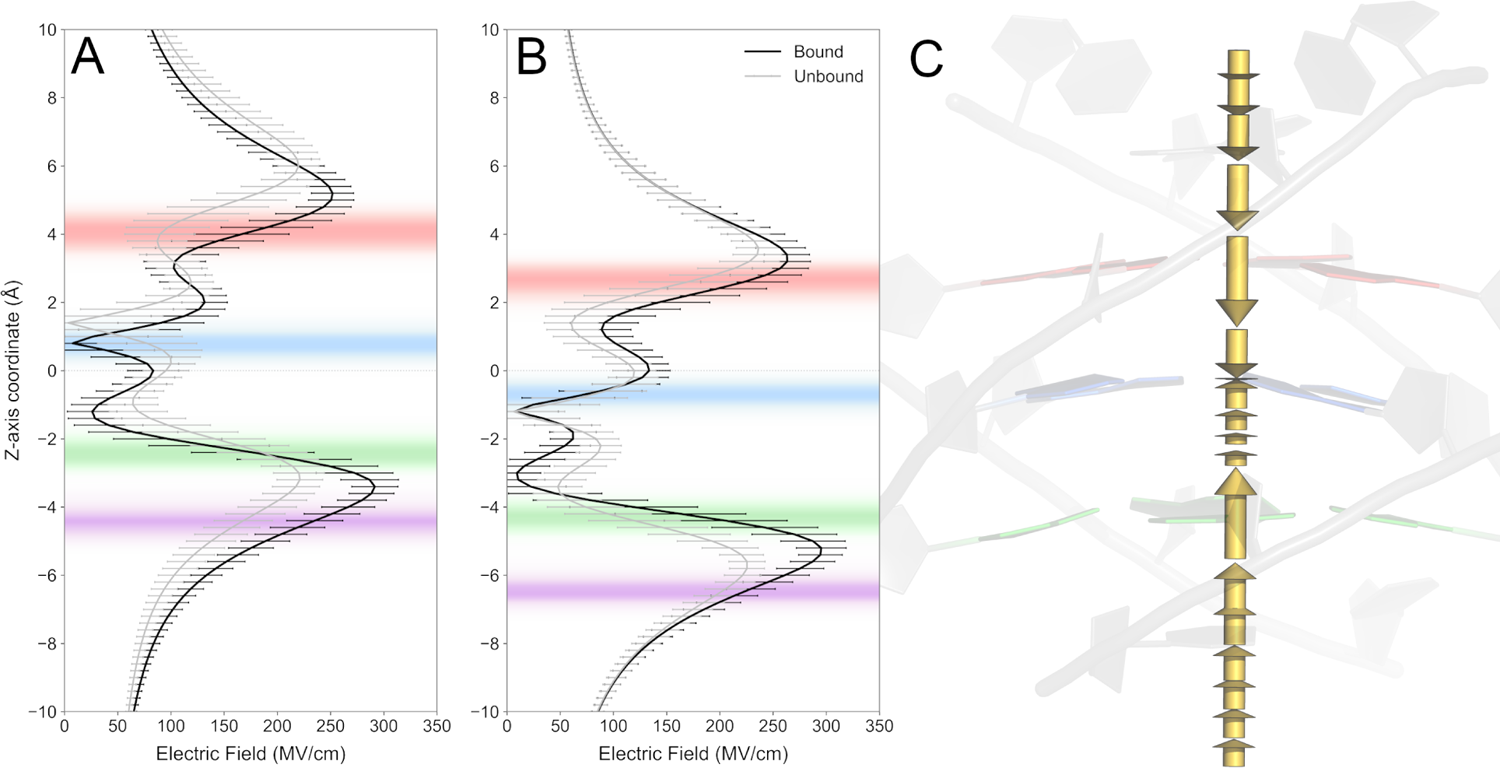
Values of *|E^→^_Z_|* along the O6-core axis for the (A) telomeric and (B) TERRA GQs for the bound states (black line). *E^→^_Z_*profiles of the unbound state are shown in gray for reference. The color and rendering scheme is the same as in Figure 6. Field vectors are shown in (C), with vector lengths proportional to the field magnitude along the O6-core.

Despite the same field values at the bound states, it is important to note that a *≈*230 MV/cm field for both GQs represents an increase of *≈*70 MV/cm for the telomeric GQ and only 30 MV/cm for the TERRA GQ, suggesting that the tetrad dynamics of TERRA contribute to a strong electric field prior to ion binding events and, therefore, tetrad fluctuations are unlikely to impact the strength of *E^→^_Z_*at S3. In contrast, similar fluctuations are potentially more impactful for the telomeric GQ, which may explain the faster rate of K^+^ unbinding events for the telomeric GQ observed in unbiased simulations.^25^

Another important feature to consider when discussing these electric fields is the impact of explicit electronic polarization. To quantify the extent to which polarization contributes to the observed electric fields, we recalculated all *E^→^_Z_*profiles along the O6-core axis using topologies in which the charges of the Drude oscillators were merged into those of their parent atoms. By doing so, we canceled the induced polarization response and analyzed only the geometric contribution of the residues to *E^→^_Z_*.

As shown in Figures 8 and 9, the induced polarization is a major component to the fields acting at S3 for both GQs. In the unbound state, the explicit polarization contributes *≈*60 MV/cm and *≈*75 MV/cm to *E^→^_Z_*exerted at S3 by the telomeric and TERRA GQs, respectively. In the bound states, these polarization contributions increased to *≈*95 MV/cm for both systems. Overall, the absence of explicit polarization did not shift the *E^→^_Z_*profiles in any direction and only impacted the field magnitudes. These results not only suggest that accounting for explicit polarization response is critical when modeling GQs, but they also support our hypothesis that shifts in these field profiles are caused by the tetrad dynamics almost exclusively.

**Figure 8:**
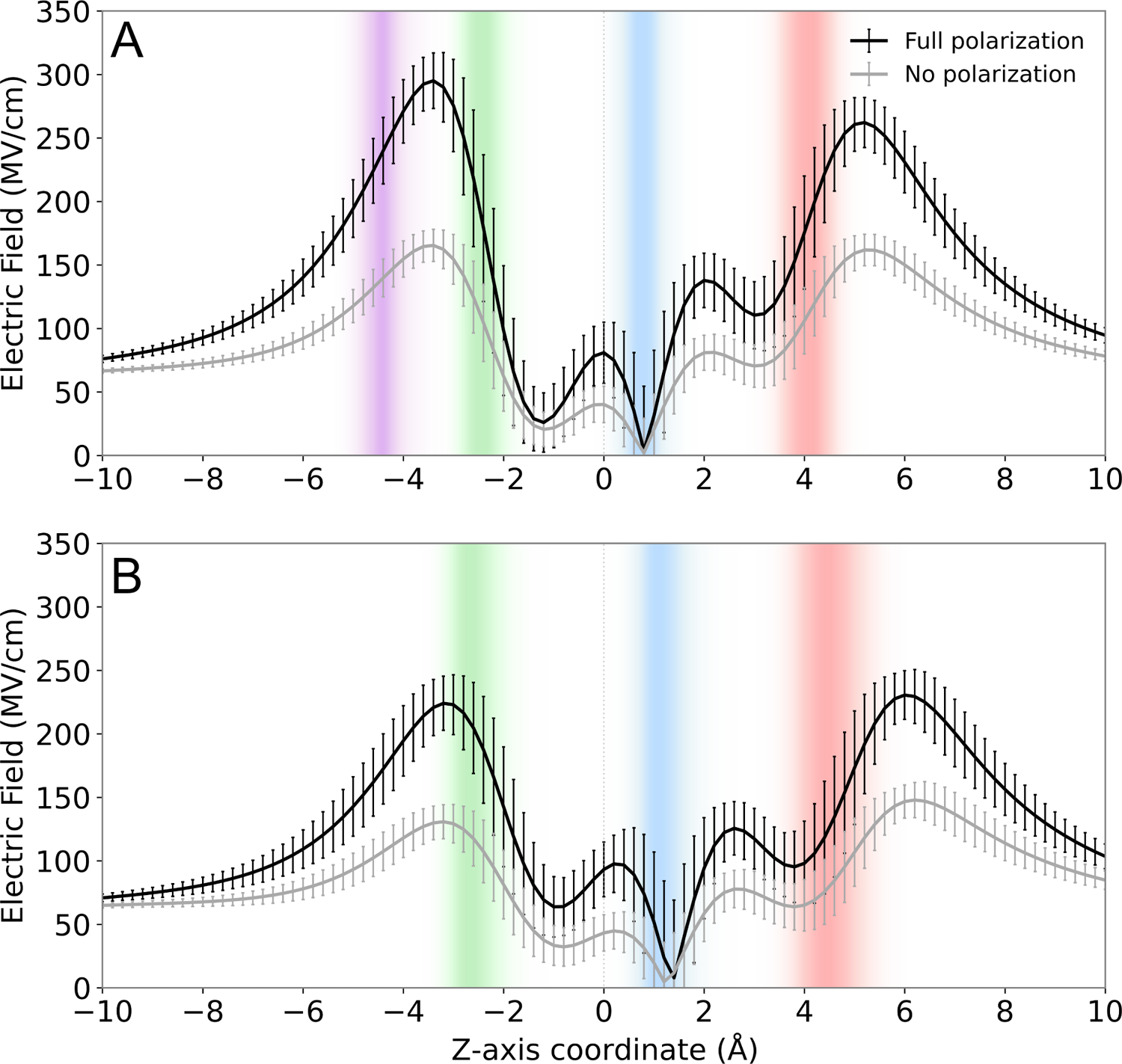
Z-axis electric field component exerted by the Telomeric GQ across its O6-core axis for its bound (A) and unbound (B) state considering the presence and absence of induced polarization (black and gray lines, respectively). Z-axis location of T1, T2 and T3 guanine O6 atoms in the unbound state are shown as distributions in red, blue and green, respectively.

**Figure 9:**
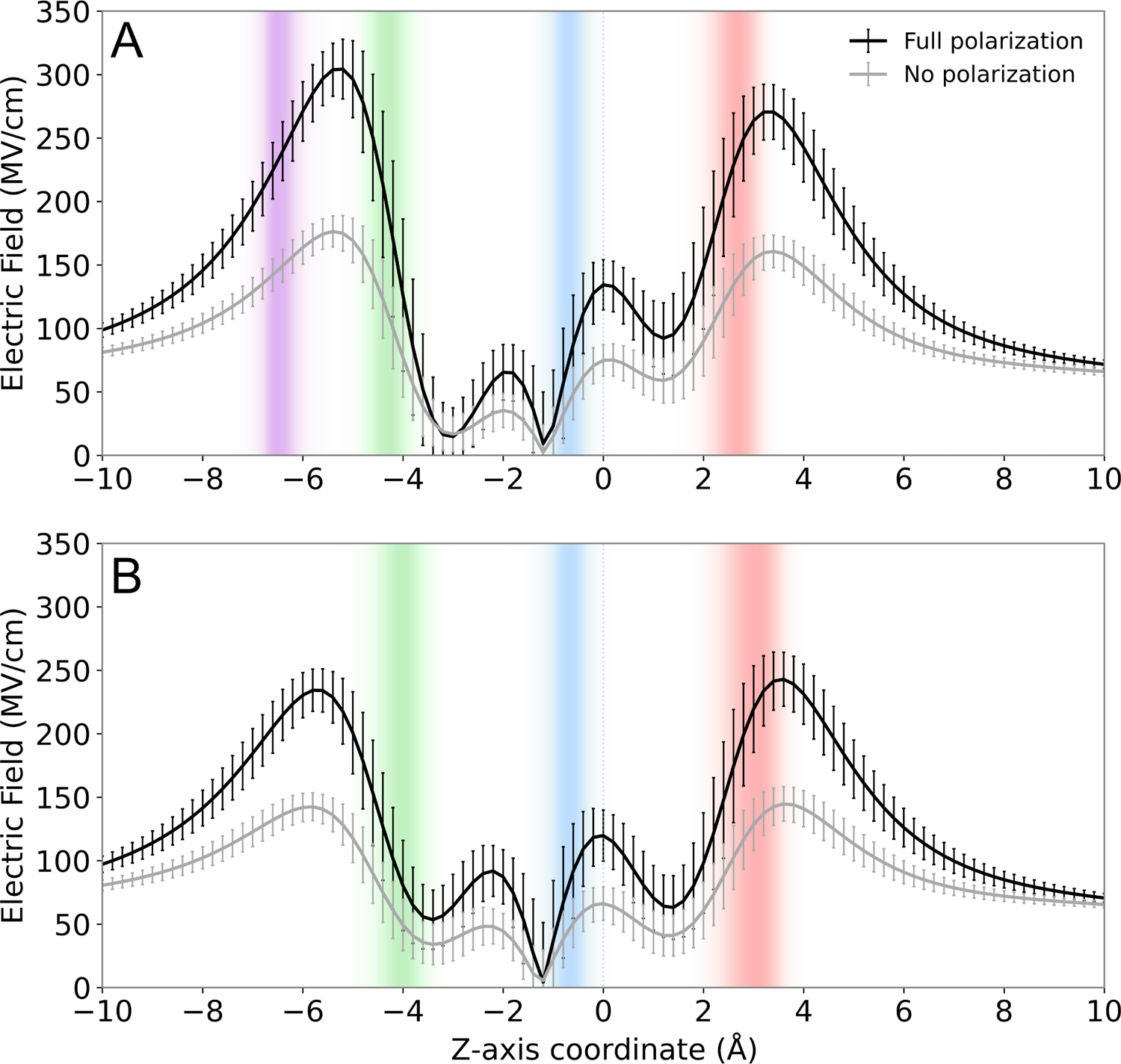
Z-axis electric field component exerted by the TERRA GQ across its O6-core axis for its bound (A) and unbound (B) state considering the presence and absence of induced polarization (black and gray lines, respectively). Z-axis location of T1, T2 and T3 guanine O6 atoms in the unbound state are shown as distributions in red, blue and green, respectively.

Importantly, *E^→^*profiles along the O6-core axis were calculated while neglecting the contributions of residues 1, 2 and 12, and the coordinated ions K1 and K2 to the field exerted at the binding K^+^ ion. Such a simplification was necessary to accurately calculate the field along the O6-axis and to understand the contribution of the tetrad dynamics to the electric field profiles. To understand the actual electric field sensed by ions binding at S3 (*E^→K^*^+^), we used TUPÃ to calculate *E^→K^*^+^ considering the contributions of the entire GQ and the coordinated ions K1 and K2 (Figure 10).

**Figure 10:**
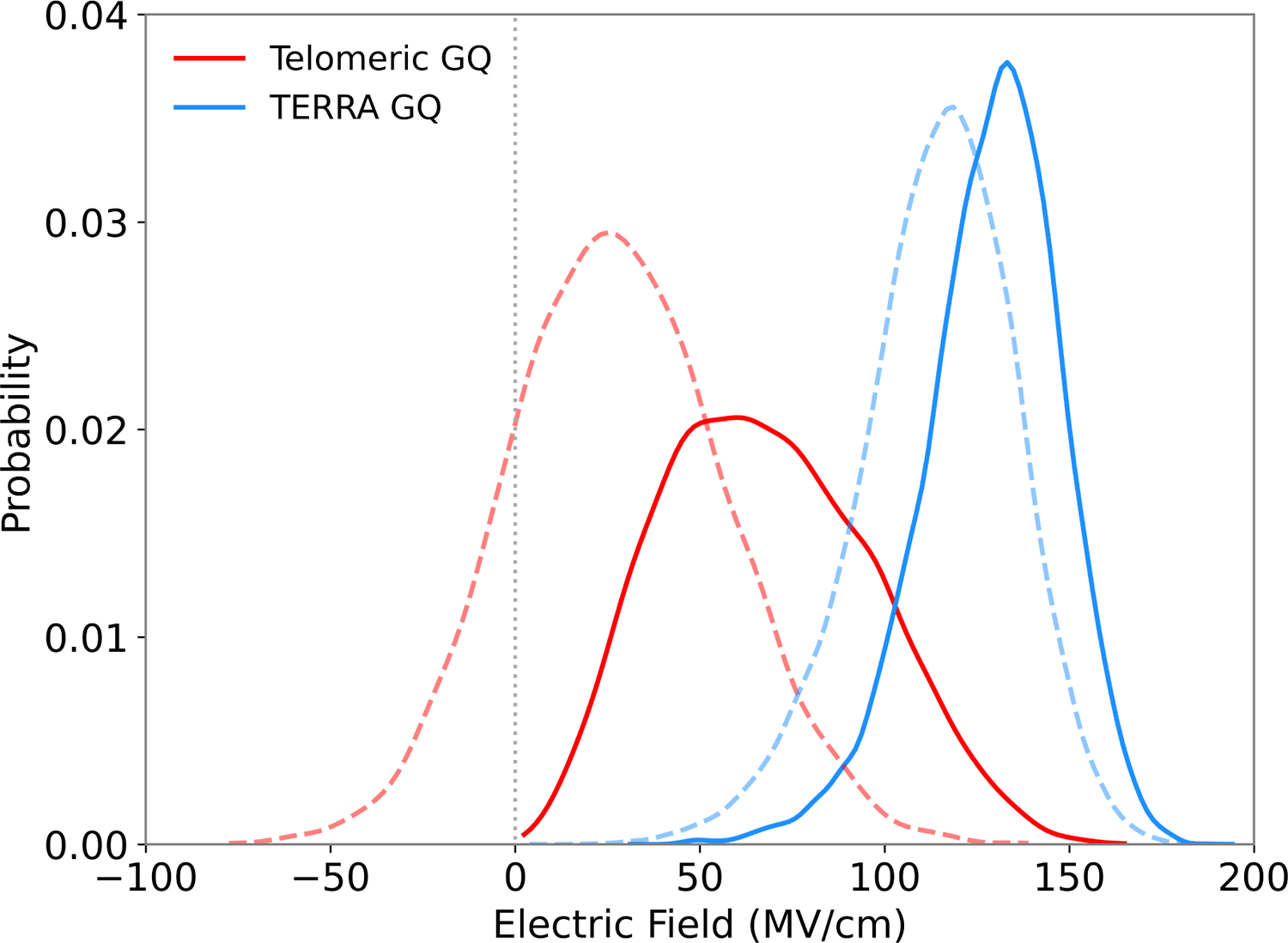
Distributions of *|E^→ K^*^+^ | (full lines) and their corresponding *z*-axis components *E^→ K^*^+^ (dashed lines) for telomeric and TERRA GQs.

Overall, the K^+^ ions binding at S3 feel a weaker field exerted by the telomeric GQ than that of the TERRA GQ at the same site. On average, *|E^→K^*^+^ *|* of the telomeric GQ was 67 *±* 28 MV/cm, whereas for the TERRA GQ, *|E^→K^*^+^ *|* was 129 *±* 18 MV/cm, suggesting that ions are able to bind more strongly to TERRA GQ S3 than the telomeric GQ S3. Such a large difference is in line with our metadynamics calculations and the binding patterns observed in the unbiased simulations reported previously.^25^ Moreover, by decomposing the *E^→K^*^+^into its *z*-axis component (*E^→K^*^+^), it is clear that while the field exerted by the telomeric GQ sampled negative values (repelling the ion towards the bulk region and thereby facilitating dissociation), *E^→K^*^+^ exerted by the TERRA GQ was always positive (attracting the ion towards S3 and favoring binding).

The observed electronic phenomenon in the telomeric GQ is linked to T3 electronic polarization. We calculated the average induced dipole moment of all T3 bases in the bound and unbound states and investigated the *x, y* and *z*-axis dipole components. As expected, both GQs showed little to no T3 electronic polarization along the *x* and *y*-axes upon K^+^ binding, reflecting the symmetrical nature of a GQ tetrad. Interestingly, the average *z*-axis component of T3 in telomeric GQ in the unbound state was −2.8 *±* 0.9 D, with a negative sign reflecting its dipole moment oriented towards S3 and thus, reflecting an accumulation of negative electron density at the outer face of T3. In comparison, the average *z*-axis component of T3 in TERRA GQ was 0.3 *±* 0.9 D.

Upon K^+^ binding at S3, the average *z*-axis component of T3 in telomeric GQ was 2.0 *±* 0.8 D, unexpectedly inverting the dipole moment orientation towards K2. At the same time, the K^+^ ion at S3 manifested an average *z*-axis dipole component of 0.13 *±* 0.07 D, reflecting an accumulation of the ion’s electron density at the outer face of T3. For the TERRA GQ, the average *z*-axis component of T3 was only 0.5 *±* 0.9 D, while the K^+^ ion binding at S3 had an average *z*-axis dipole component of 0.0 *±* 0.06 D. A complete and pronounced reorientation of T3 dipole moment was only observed for telomeric GQ, as shown in Figure 11A.

**Figure 11:**
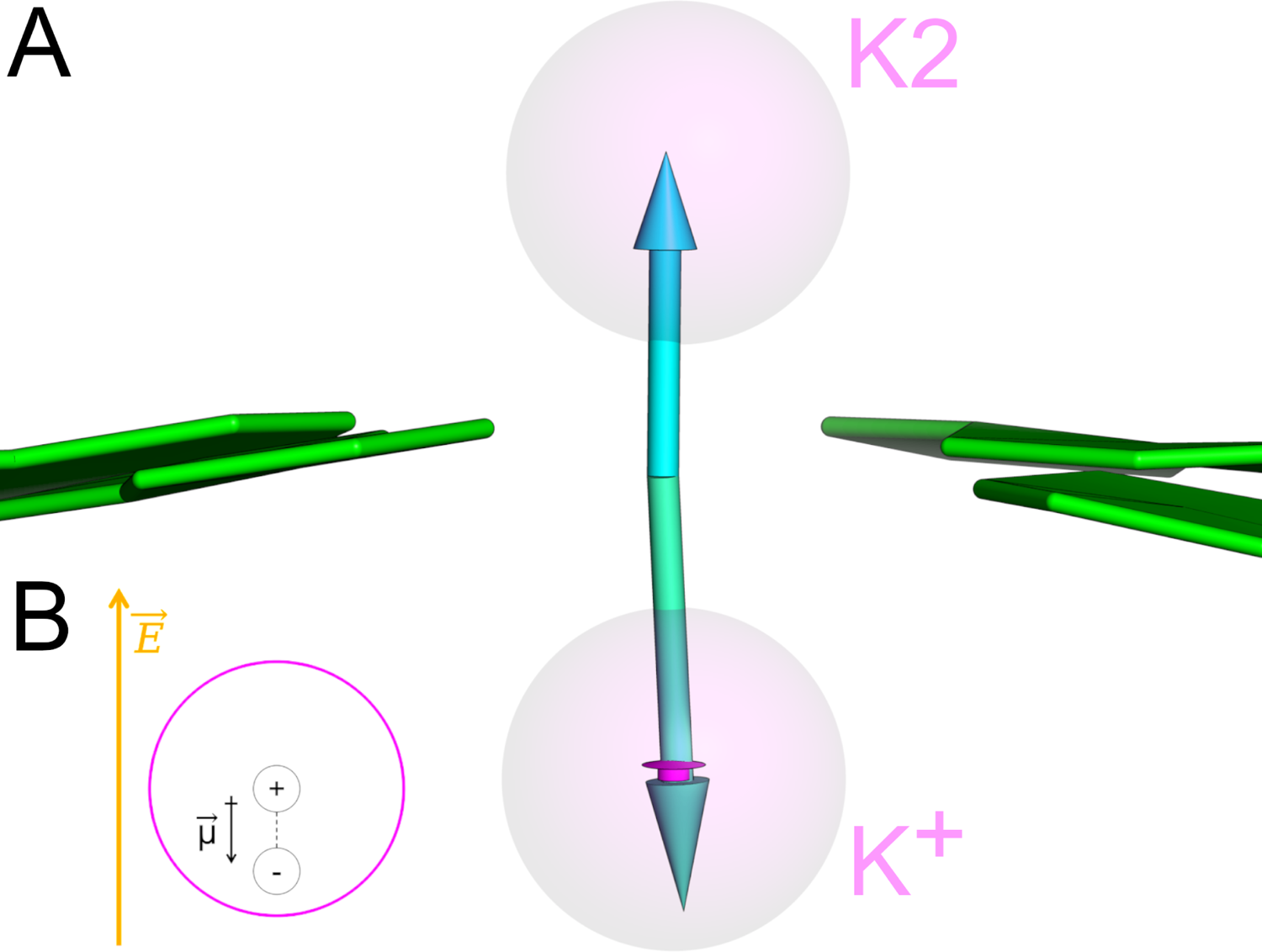
(A) Orientation of the T3 dipole moments of the telomeric GQ in the unbound state (greencyan) and bound state (cyan). The dipole moment of the binding K^+^ ion is shown in magenta. (B) Scheme of a Drude oscillator (negative sphere) displacing inside its parent atom (magenta) under the influence of an electric field (orange).

As shown in Figure 11B, the usual response of a negatively charged Drude oscillator at S3 would be to displace in opposition to the electric field, yielding a dipole moment oriented in the other direction. However, we observed the contrary in both GQ systems, but more much more pronounced in the telomeric GQ. This observation suggests that the multibody polarization effects arising from the local microenvironment at the T3/S3 interface somewhat oppose the effects of global *E^→^_Z_*acting on K^+^, explaining the weaker ion binding reported for the telomeric GQ.

We acknowledge that experimentally testing the electronic properties of binding ions captured by our model and how that could impact binding affinity is challenging, but the principles shown here should be applicable to GQ ligands binding to tetrad interfaces, such as N-methyl mesoporphyrins,^10,37^ which in turn could be further studied in terms of TERRA and telomeric GQ binding affinities and T3/ligand interfaces properties. Understanding the fine balance between electronic properties and conformational dynamics of these two GQs could provide valuable insights for drug design efforts aiming for selectivity.

Lastly, the electric field profiles reported here could also explain the heme peroxidase/peroxygenase catalytic activity observed previously for some GQs.^38–40^ In these GQs, the heme substrate stacks on tetrad surfaces such that Fe^2+^ occupies sites like S1 and S3, where we show the field magnitudes are greatest. Studies of electric fields driving enzyme catalysis report field magnitudes ranging from *≈*40 to 140 MV/cm,^41–43^ thus the strong and oriented fields reported here could explain how some GQs participate in catalysis.

## Conclusion

By employing polarizable MD simulations in conjunction with enhanced sampling methods, we have extended the characterization of DNA and RNA GQ structures with the same overall architecture. Building upon the previous reports showing that telomeric and TERRA GQs behave differently, here we describe the relationship among tetrad dynamics, influence of ion binding, and intrinsic electric fields acting in these systems.

We characterized tetrad base tilting motions in both GQs, showing how each GQ tetrad core has intrinsic tilting directions that are modulated by K^+^ binding close to T3. We also demonstrated how (deoxy)ribose puckering angles of each tetrad nucleotide can drastically differ in telomeric and TERRA GQs, although we could not confirm whether or not the sugar conformation and base tilting motions were linked.

Moreover, we characterized the polarization response of the tetrad guanine bases upon K^+^ binding, showing the impact of including explicit electronic polarization terms in the force field via interaction energy calculations. Overall, the TERRA GQ interacted more strongly with K^+^ ions than the telomeric GQ. These results were further confirmed by well-tempered cylinder-based metadynamics simulations, from which we calculated absolute ion binding free energies. Lastly, we investigated how these GQs organize electric fields around themselves to attract ions. We found that these fields are responsible for the focusing of binding sites close to exposed tetrads and that they are impacted by tetrad dynamics and K^+^ binding. In line with the free energy calculations, we observed a stronger field exerted by TERRA GQ at the K^+^ ion bound to binding site S3 in comparison to that of the telomeric GQ. Moreover, we identified a fine balance between electric field and dipole moment orientation at T3/S3 interface that explains the weaker binding of K^+^ ions to the telomeric GQ.

Together, these results quantify differences between two GQs with very similar architectures, attesting to the structural and electronic diversity of GQs and suggesting important intrinsic differences between DNA and RNA. Moreover, the metadynamics protocol employed here in conjunction with the electric field analyses can be further extended to more complex ligands and GQ structures, guiding future drug design efforts and investigations into interactions with other cellular biomolecules like proteins.

## Methods

### Tetrad base tilting

To calculate tilting angle of each tetrad guanine in relation to its tetrad plane, we defined a out-of-plane angle by using a pseudo improper dihedral scheme. The tetrad plane was defined by the C1’ atom of the guanine under analysis (G) and the two C1’ atoms of its neighboring guanines (G+ and G-). The O6 atom of G was used to define the out-of-plane angle (see Figure S6).

### Repulsive potential simulations

To understand the relationship between the structural dynamics of the GQs and the binding of bulk K^+^ ions to the tetrad surfaces, a repulsive potential was applied to all bulk ions to prevent ion binding at S1 and S3. Doing so allowed us to decouple ion binding from intrinsic dynamics. We used the K1 and K2 positions as references to define two radial cutoff distances (*D_K_*_1_ and *D_K_*_2_, respectively) between the bound ions and the closest bulk ion (Figure S1).

To do so, we used the PLUMED plugin^44,45^ in conjunction with OpenMM, applying *LOWER_WALLS* setups (one for *D_K_*_1_ and another one for *D_K_*_2_) as shown in equation 1. These biased simulations were performed in 3 replicates of 1 *µ*s each for both the telomeric and TERRA GQs. For consistency, the remaining simulation parameters were the same as the corresponding unbiased simulations.^25^

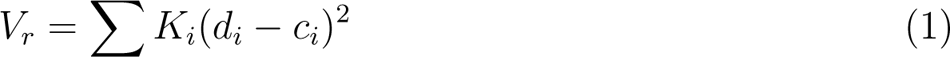

In our scheme, *d_i_*is the minimum distance of any bulk ion to the closest bound K^+^ ion (either K1 or K2), *c_i_*is the cutoff distance (set to 5.5 Å), and *K_i_* was set to 2000 kJ/mol for both potentials.

Despite its simplicity, this approach was proved useful since our 5.5 Å cutoff only encompassed tetrad 1 and tetrad 3 binding sites, allowing ion binding to other GQ sites (see Figure S1). Furthermore, the distance between tetrad 1 and tetrad 3 binding sites is greater than 5.5 Å, so only one of the two repulsive forces was exerted on bulk ions eventually crossing the cutoff distance. For reproduction purposes, PLUMED scripts for repulsive potential and well-tempered cylinder metadynamics simulations are available at the PLUMED-NEST consortium (https://www.plumed-nest.org/eggs/23/025/).

### Well-tempered cylinder metadynamics

To probe K^+^ binding free energy to each GQ, we performed a volume-based well-tempered metadynamics (WT-MetaD) largely inspired by the work of Capelli et al. ^35^. While the authors used spherical coordinates as collective variables (CVs), here we used cylinder coordinates (radial distance *ρ*, azimuth *φ*, and height *ζ*) to define a cylindrical volume that encompasses the tetrad binding site of interest and the bulk region. One K^+^ ion was defined as ligand and had its sampling confined to the cylindrical volume. To avoid competition between ligand and bulk ions, we also employed the repulsive potential scheme described above such that the only ion able to bind to the GQ tetrads was the ligand.

The overall structure of the GQs was centered at the Cartesian origin such that the tetrad planes were parallel to the XY plane and the O6-core axis was aligned to the *z*-axis. The radius of the cylinder *ρ* was limited to 7 Å, which is about the maximum radius of the tetrads in relation to the O6-core axis. To define the height of the cylinder, the T1-T2 and T2-T3 centers-of-mass were defined as COM_K1_ and COM_K2_, respectively, and used as reference points to calculate the distance to the ligand position. For S1, the distance was calculated between COM_K1_ and the ligand and COM_K2_ was used for S3. The *z*-axis component of this distance was used as cylinder height *ζ*.

WT-MetaD simulations were performed for 500 ns by biasing all three cylindrical coordinates mentioned above as CVs. Gaussian hills with a height of 1.0 kJ/mol were deposited every 0.5 ps, the bias factor *λ* was set to 5 and sigmas for *ρ*, *φ* and *ζ* were set to 0.5 Å, *π*/8 rad and 0.5 Å, respectively. The GQ core was realigned to the initial coordinates every step to maintain a fixed reference for the CVs.

Reweighting procedures used instantaneous values of the bias calculated during simulation and absolute Cartesian coordinates of the ligand calculated via CHARMM^46^ after reorientation of the trajectory frames using tetrad 2 C1’ atoms as reference for centering the GQ at Cartesian origin. As before, the GQ O6-core axis was aligned to the *z*-axis.

### TUPÃ calculations

To calculate electric fields acting in our simulations, we used the TUPÃ algorithm,^36^ which is compatible with Drude polarizable model. While the TUPÃ parameters for each type of calculation are provided in the Open Science Framework at https://osf.io/2pfj8/, below we overview some aspects of the setup.

#### Electric field across O6-core axis

The O6-core axis, in GQs encompasses both coordinated K^+^ ions and binding sites S1 and S3. To measure the electric field acting across this axis, both the telomeric and TERRA GQ trajectories were oriented so the Z-axis was aligned to the GQs O6-core axis. Also, the T2 C1’ atoms were used as reference to center the GQ at cartesian origin.

Using COORDINATE mode in TUPÃ, the time series of electric field magnitudes and directions were calculated at coordinates [0,0,*z*], where −10 *≤ z ≤* 10 Å with a step of 0.2 Å. To understand the intrinsic field exerted by the GQ itself, residues 3 to 11 were selected as *environment set* (i.e., the atoms exerting the electric field) and no solvent molecules (water or ions) or coordinated K^+^ ions were considered for this calculation. The average and standard deviation of each time series were calculated via Numpy.^47^

#### Electric field at K^+^ ligand

First, we extracted only the frames in which K^+^ ions were occupying the binding site S3 from the unbiased simulation trajectories. Then, using TUPÃ’s ATOM mode, we selected the entire GQ and the coordinated ions K1 and K2 to act as the *environment set*. Probe atoms were selected by using MDAnalysis^48,49^ dynamical selection syntax to select any K^+^ atoms within 1 Å from S3. To guarantee accuracy, we manually confirmed that such selection would capture only one binding ion. The average and standard deviation of each electric field time series were calculated via Numpy^47^

## Acknowledgement

This work was supported by the National Institutes of Health (grant R35GM133754 to JAL) and USDA-NIFA (project VA-160092). The authors thank Virginia Tech Advanced Research Computing for computing time and resources.

## Supporting Information Available

Supporting Information is available free of charge at https://pubs.acs.org. Schematic for the repulsive potential scheme, x- and y-components of electric fields along the tetrad axis in telomeric and TERRA GQs, and tilting angle calculation scheme.

## TOC Graphic

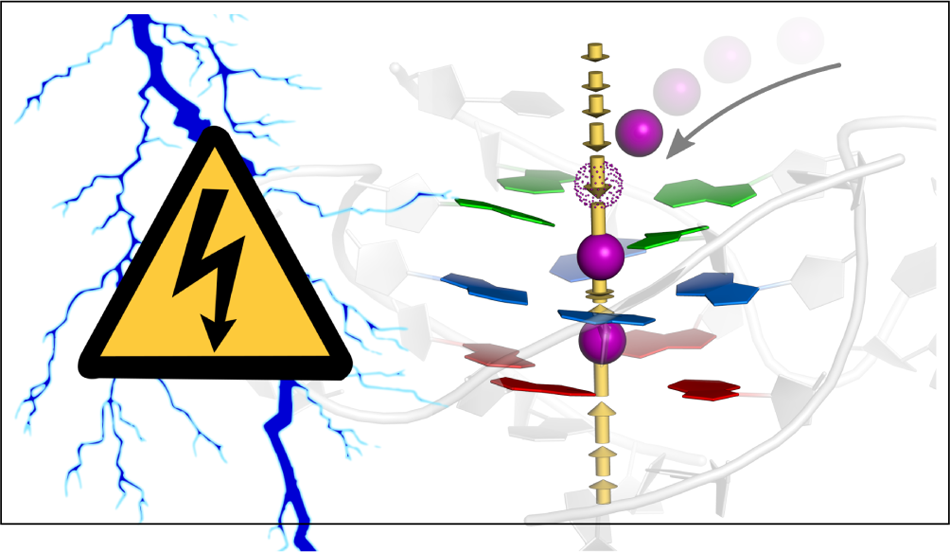

## References

1. Duxin, J. P.; Dao, B.; Martinsson, P.; Rajala, N.; Guittat, L.; Campbell, J. L.; Spelbrink, J. N.; Stewart, S. A. Human Dna2 Is a Nuclear and Mitochondrial DNA Maintenance Protein. Molecular and Cellular Biology 2009, 29, 4274–4282.

2. Wu, Y.; Shin-ya, K.; Brosh, R. M. FANCJ Helicase Defective in Fanconia Anemia and Breast Cancer Unwinds G-Quadruplex DNA To Defend Genomic Stability. Molecular and Cellular Biology 2008, 28, 4116–4128.

3. Besnard, E.; Babled, A.; Lapasset, L.; Milhavet, O.; Parrinello, H.; Dantec, C.; Marin, J.-M.; Lemaitre, J.-M. Unraveling cell type–specific and reprogrammable human replication origin signatures associated with G-quadruplex consensus motifs. Nature Structural & Molecular Biology 2012, 19, 837–844.

4. Cayrou, C.; Coulombe, P.; Vigneron, A.; Stanojcic, S.; Ganier, O.; Peiffer, I.; Rivals, E.; Puy, A.; Laurent-Chabalier, S.; Desprat, R.; Méchali, M. Genome-scale analysis of metazoan replication origins reveals their organization in specific but flexible sites defined by conserved features. Genome Research 2011, 21, 1438–1449.

5. Moye, A. L.; Porter, K. C.; Cohen, S. B.; Phan, T.; Zyner, K. G.; Sasaki, N.; Lovrecz, G. O.; Beck, J. L.; Bryan, T. M. Telomeric G-quadruplexes are a substrate and site of localization for human telomerase. Nature Communications 2015, 6, 7643, Number: 1 Publisher: Nature Publishing Group.

6. Bugaut, A.; Balasubramanian, S. 5’-UTR RNA G-quadruplexes: translation regulation and targeting. Nucleic Acids Research 2012, 40, 4727–4741.

7. Millevoi, S.; Moine, H.; Vagner, S. G-quadruplexes in RNA biology. *Wiley interdisciplinary reviews*. RNA 2012, 3, 495–507.

8. Lim, K. W.; Amrane, S.; Bouaziz, S.; Xu, W.; Mu, Y.; Patel, D. J.; Luu, K. N.; Phan, A. T. Structure of the Human Telomere in K^+^ Solution: A Stable Basket-Type G-Quadruplex with Only Two G-Tetrad Layers. Journal of the American Chemical Society 2009, 131, 4301–4309, PMID: 19271707.

9. Parkinson, G. N.; Lee, M. P. H.; Neidle, S. Crystal structure of parallel quadruplexes from human telomeric DNA. 417, 876–880, Number: 6891 Publisher: Nature Publishing Group.

10. Han, F. X.; Wheelhouse, R. T.; Hurley, L. H. Interactions of TMPyP4 and TMPyP2 with Quadruplex DNA. Structural Basis for the Differential Effects on Telomerase Inhibition. Journal of the American Chemical Society 1999, 121, 3561–3570, Publisher: American Chemical Society.

11. Kim, M.-Y.; Vankayalapati, H.; Shin-Ya, K.; Wierzba, K.; Hurley, L. H. Telomestatin, a potent telomerase inhibitor that interacts quite specifically with the human telomeric intramolecular g-quadruplex. Journal of the American Chemical Society 2002, 124, 2098–2099.

12. Cusanelli, E.; Chartrand, P. Telomeric repeat-containing RNA TERRA: a noncoding RNA connecting telomere biology to genome integrity. Frontiers in Genetics 2015, 6.

13. Collie, G. W.; Haider, S. M.; Neidle, S.; Parkinson, G. N. A crystallographic and modelling study of a human telomeric RNA (TERRA) quadruplex. Nucleic Acids Research 2010, 38, 5569–5580.

14. Mei, Y.; Deng, Z.; Vladimirova, O.; Gulve, N.; Johnson, F. B.; Drosopoulos, W. C.; Schildkraut, C. L.; Lieberman, P. M. TERRA G-quadruplex RNA interaction with TRF2 GAR domain is required for telomere integrity. Scientific Reports 2021, 11, 3509.

15. Balasubramanian, S.; Hurley, L. H.; Neidle, S. Targeting G-quadruplexes in gene promoters: a novel anticancer strategy? Nature reviews. Drug discovery 2011, 10, 261–275.

16. Šponer, J.; Špačková, N. Molecular dynamics simulations and their application to four-stranded DNA. Methods 2007, 43, 278–290.

17. Li, N.; Gao, Y.; Qiu, F.; Zhu, T. Benchmark Force Fields for the Molecular Dynamic Simulation of G-Quadruplexes. Molecules 2021, 26, 5379, Number: 17 Publisher: Multidisciplinary Digital Publishing Institute.

18. Green, A. T.; Pickard, A. J.; Li, R.; MacKerell, A. D. J.; Bierbach, U.; Cho, S. S. Computational and Experimental Characterization of rDNA and rRNA G-Quadruplexes. The Journal of Physical Chemistry B 2022, 126, 609–619, Publisher: American Chemical Society.

19. Gkionis, K.; Kruse, H.; Platts, J. A.; Mládek, A.; Koča, J.; Šponer, J. Ion Binding to Quadruplex DNA Stems. Comparison of MM and QM Descriptions Reveals Sizable Polarization Effects Not Included in Contemporary Simulations. Journal of Chemical Theory and Computation 2014, 10, 1326–1340, Publisher: American Chemical Society.

20. Salsbury, A. M.; Lemkul, J. A. Molecular Dynamics Simulations of the c-kit1 Promoter G-Quadruplex: Importance of Electronic Polarization on Stability and Cooperative Ion Binding. The Journal of Physical Chemistry B 2019, 123, 148–159.

21. Salsbury, A. M.; Dean, T. J.; Lemkul, J. A. Polarizable Molecular Dynamics Simulations of Two c-kit Oncogene Promoter G-Quadruplexes: Effect of Primary and Secondary Structure on Loop and Ion Sampling. Journal of Chemical Theory and Computation 2020, 16, 3430–3444, Publisher: American Chemical Society.

22. Salsbury, A. M.; Lemkul, J. A. Cation competition and recruitment around the c-kit1 G-quadruplex using polarizable simulations. Biophysical Journal 2021, 120, 2249–2261.

23. Salsbury, A. M.; Michel, H. M.; Lemkul, J. A. Ion-Dependent Conformational Plasticity of Telomeric G-Hairpins and G-Quadruplexes. ACS Omega 2022, 7, 23368–23379, Publisher: American Chemical Society.

24. Ratnasinghe, B. D.; Salsbury, A. M.; Lemkul, J. A. Ion Binding Properties and Dynamics of the *bcl*2 G-Quadruplex Using a Polarizable Force Field. Journal of Chemical Information and Modeling 2020, 60, 6476–6488.

25. Lemkul, J. A. Same fold, different properties: polarizable molecular dynamics simulations of telomeric and TERRA G-quadruplexes. Nucleic Acids Research 2020, 48, 561–575.

26. Lemkul, J. A.; MacKerell, A. D. J. Polarizable Force Field for DNA Based on the Classical Drude Oscillator: I. Refinement Using Quantum Mechanical Base Stacking and Conformational Energetics. Journal of Chemical Theory and Computation 2017, 13, 2053–2071, Publisher: American Chemical Society.

27. Lemkul, J. A.; MacKerell, A. D. J. Polarizable Force Field for DNA Based on the Classical Drude Oscillator: II. Microsecond Molecular Dynamics Simulations of Duplex DNA. Journal of Chemical Theory and Computation 2017, 13, 2072–2085, Publisher: American Chemical Society.

28. Lemkul, J. A.; MacKerell Jr., A. D. Polarizable force field for RNA based on the classical drude oscillator. Journal of Computational Chemistry 2018, 39, 2624–2646, _eprint: https://onlinelibrary.wiley.com/doi/pdf/10.1002/jcc.25709.

29. Martadinata, H.; Phan, A. T. Structure of Propeller-Type Parallel-Stranded RNA G-Quadruplexes, Formed by Human Telomeric RNA Sequences in K+ Solution. Journal of the American Chemical Society 2009, 131, 2570–2578, Publisher: American Chemical Society.

30. Altona, C.; Sundaralingam, M. Conformational analysis of the sugar ring in nucleosides and nucleotides. New description using the concept of pseudorotation. Journal of the American Chemical Society 1972, 94, 8205–8212, Publisher: American Chemical Society.

31. Cheong, C.; Moore, P. B. Solution structure of an unusually stable RNA tetraplex containing G- and U-quartet structures. Biochemistry 1992, 31, 8406–8414, Publisher: American Chemical Society.

32. Deng, J.; Xiong, Y.; Sundaralingam, M. X-ray analysis of an RNA tetraplex (UGGGGU)4 with divalent Sr2+ ions at subatomic resolution (0.61 Å). Proceedings of the National Academy of Sciences 2001, 98, 13665–13670, Publisher: Proceedings of the National Academy of Sciences.

33. Marcos-Alcalde, I.; López-Viñas, E.; Gómez-Puertas, P. MEPSAnd: minimum energy path surface analysis over n-dimensional surfaces. Bioinformatics 2020, 36, 956–958.

34. Allen, T. W.; Andersen, O. S.; Roux, B. Energetics of ion conduction through the gramicidin channel. Proceedings of the National Academy of Sciences 2004, 101, 117–122, Publisher: Proceedings of the National Academy of Sciences.

35. Capelli, R.; Carloni, P.; Parrinello, M. Exhaustive Search of Ligand Binding Pathways via Volume-Based Metadynamics. The Journal of Physical Chemistry Letters 2019, 10, 3495–3499, Publisher: American Chemical Society.

36. Polêto, M. D.; Lemkul, J. A. TUPÃ: Electric field analyses for molecular simulations. Journal of Computational Chemistry 2022, 43, 1113–1119, _eprint: https://onlineli-brary.wiley.com/doi/pdf/10.1002/jcc.26873.

37. Nicoludis, J. M.; Miller, S. T.; Jeffrey, P. D.; Barrett, S. P.; Rablen, P. R.; Lawton, T. J.; Yatsunyk, L. A. Optimized End-Stacking Provides Specificity of N-Methyl Mesoporphyrin IX for Human Telomeric G-Quadruplex DNA. Journal of the American Chemical Society 2012, 134, 20446–20456, Publisher: American Chemical Society.

38. Yamamoto, Y.; Araki, H.; Shinomiya, R.; Hayasaka, K.; Nakayama, Y.; Ochi, K.; Shibata, T.; Momotake, A.; Ohyama, T.; Hagihara, M.; Hemmi, H. Structures and Catalytic Activities of Complexes between Heme and All Parallel-Stranded Monomeric G-Quadruplex DNAs. Biochemistry 2018, 57, 5938–5948, Publisher: American Chemical Society.

39. Shinomiya, R.; Katahira, Y.; Araki, H.; Shibata, T.; Momotake, A.; Yanagisawa, S.; Ogura, T.; Suzuki, A.; Neya, S.; Yamamoto, Y. Characterization of Catalytic Activities and Heme Coordination Structures of Heme–DNA Complexes Composed of Some Chemically Modified Hemes and an All Parallel-Stranded Tetrameric G-Quadruplex DNA Formed from d(TTAGGG). Biochemistry 2018, 57, 5930–5937, Publisher: American Chemical Society.

40. Ibrahim, H.; Mulyk, P.; Sen, D. DNA G-Quadruplexes Activate Heme for Robust Catalysis of Carbene Transfer Reactions. ACS Omega 2019, 4, 15280–15288, Publisher: American Chemical Society.

41. Fried, S. D.; Bagchi, S.; Boxer, S. G. Extreme electric fields power catalysis in the active site of ketosteroid isomerase. Science (New York, N.Y.) 2014, 346, 1510–1514.

42. Vaissier, V.; Sharma, S. C.; Schaettle, K.; Zhang, T.; Head-Gordon, T. Computational Optimization of Electric Fields for Improving Catalysis of a Designed Kemp Eliminase. ACS Catalysis 2018, 8, 219–227, Publisher: American Chemical Society.

43. Bradshaw, R. T.; Dziedzic, J.; Skylaris, C. K.; Essex, J. W. The Role of Electrostatics in Enzymes: Do Biomolecular Force Fields Reflect Protein Electric Fields? Journal of Chemical Information and Modeling 2020, 60, 3131–3144, Publisher: American Chemical Society.

44. Bonomi, M.; Branduardi, D.; Bussi, G.; Camilloni, C.; Provasi, D.; Raiteri, P.; Donadio, D.; Marinelli, F.; Pietrucci, F.; Broglia, R. A.; Parrinello, M. Plumed: a Portable Plugin for Free-Energy Calculations with Molecular Dynamics. Computer Physics Communications 2009, 180, 1961–1972, Publisher: Elsevier B.V.

45. Tribello, G. A.; Bonomi, M.; Branduardi, D.; Camilloni, C.; Bussi, G. PLUMED 2: New feathers for an old bird. Computer Physics Communications 2014, 185, 604–613.

46. Brooks, B. et al. CHARMM: The Biomolecular Simulation Program. Journal of computational chemistry 2009, 30, 1545–1614.

47. Harris, C. R. et al. Array programming with {NumPy}. Nature 2020, 585, 357–362, Publisher: Springer Science and Business Media {LLC}.

48. Michaud-Agrawal, N.; Denning, E. J.; Woolf, T. B.; Beckstein, O. MDAnalysis: A toolkit for the analysis of molecular dynamics simulations. Journal of Computational Chemistry 2011, 32, 2319–2327, Publisher: John Wiley & Sons, Ltd.

49. Gowers, R. J.; Linke, M.; Barnoud, J.; Reddy, T. J. E.; Melo, M. N.; Seyler, S. L.; Domański, J.; Dotson, D. L.; Buchoux, S.; Kenney, I. M.; Beckstein, O. MDAnalysis: A Python Package for the Rapid Analysis of Molecular Dynamics Simulations. Proceedings of the 15th Python in Science Conference 2016, 98–105, Publisher: SciPy.

