## Supporting Information for "Differences in Conformational Sampling and Intrinsic Electric Fields Drive Ion Binding in Telomeric and TERRA G-Quadruplexes"

### Supporting Figures

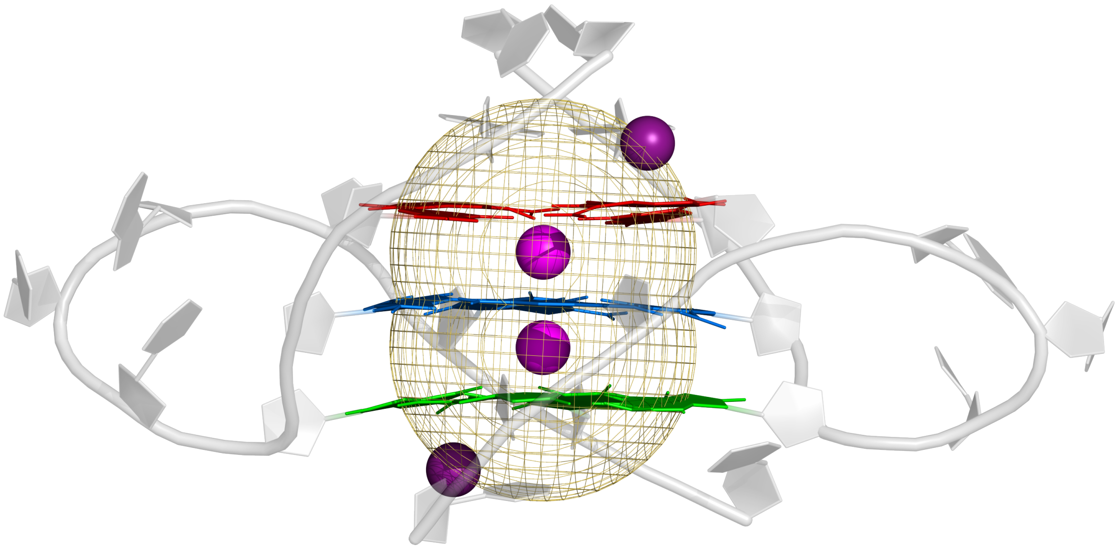

Figure S1: Repulsive potential scheme applied in our simulations to prevent ion binding at S1 and S3. A flat-bottom restraint was applied to any $K^+$ atoms within 5 Å from either the K1 or K2 atoms.

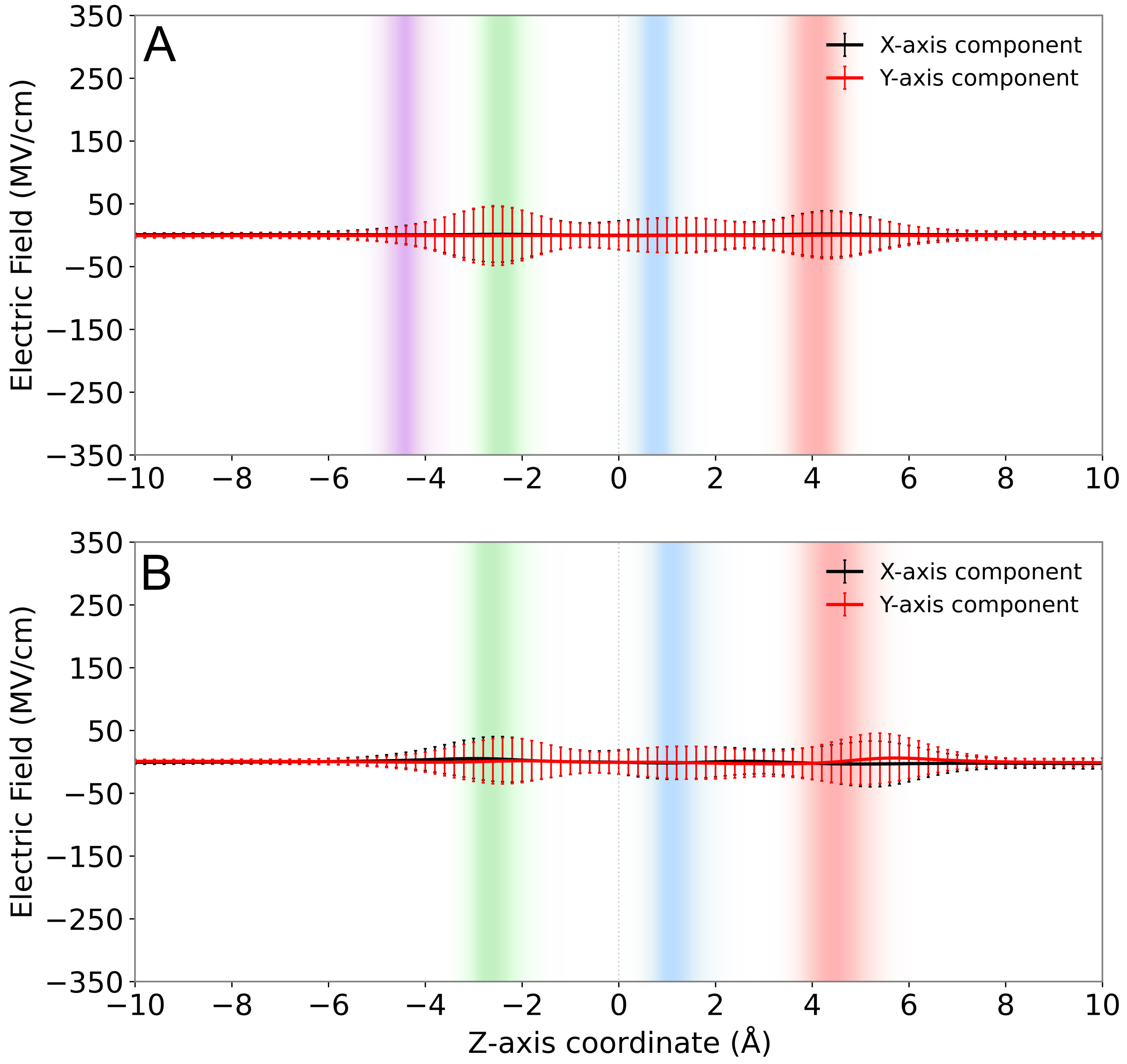


Figure S2: X-axis and Y-axis electric field components exerted by the Telomere GQ across its O6-core axis for its bound (A) and unbound (B) state. Z-axis location of T1, T2 and T3 guanine O6 atoms in the unbound state are shown as distributions in red, blue and green, respectively.

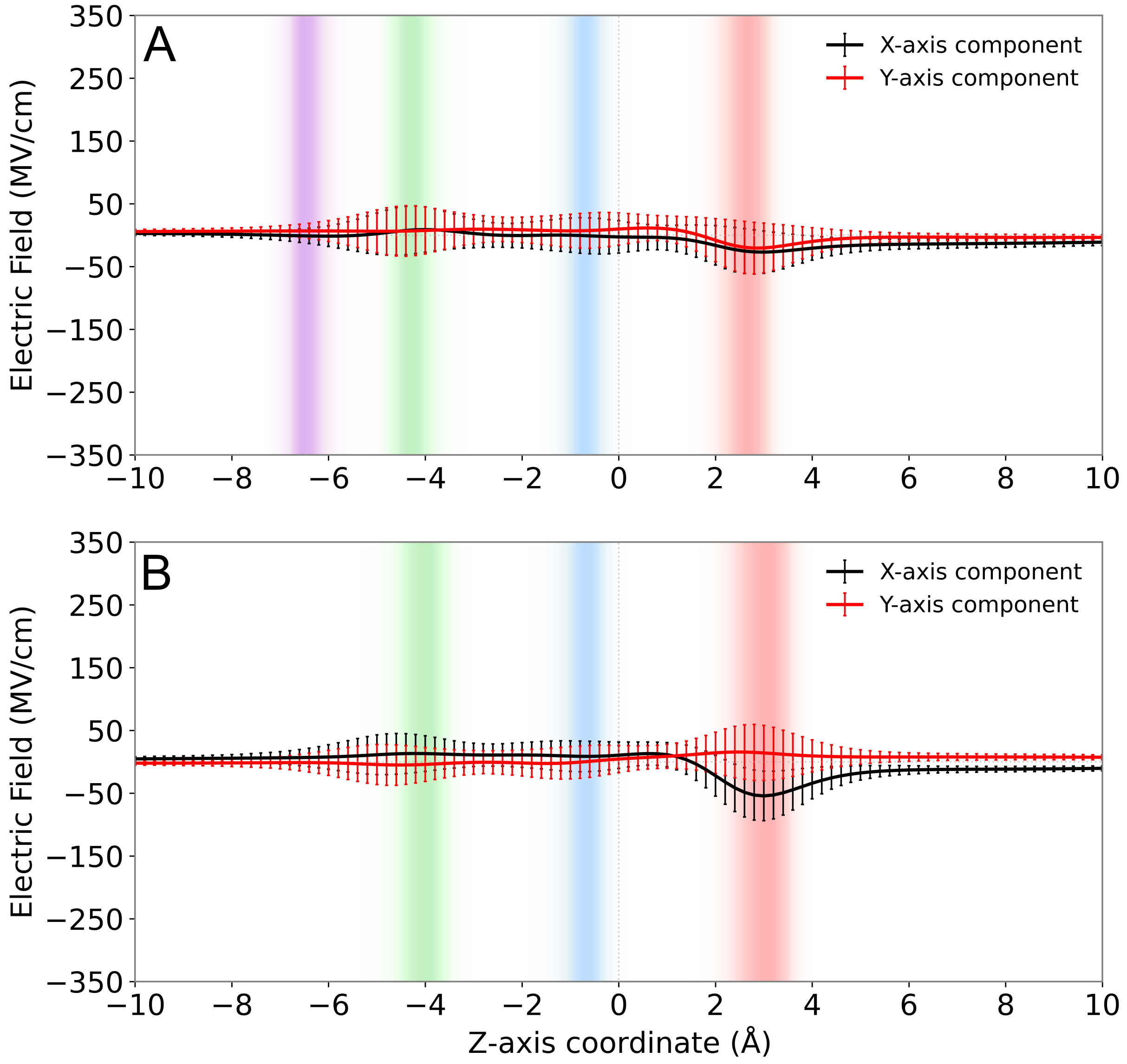


Figure S3: X-axis and Y-axis electric field components exerted by the TERRA GQ across its O6-core axis for its bound (A) and unbound (B) state. Z-axis location of T1, T2 and T3 guanine O6 atoms in the unbound state are shown as distributions in red, blue and green, respectively.

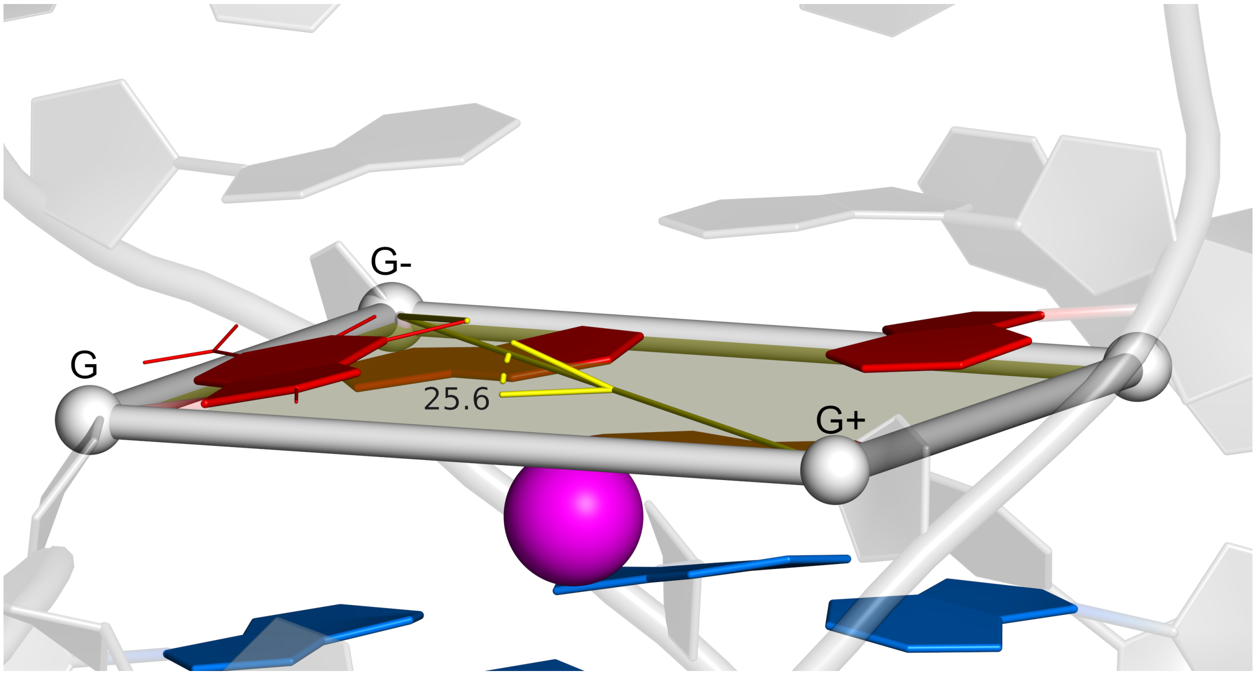


Figure S4: To calculate tilting angle of each tetrad guanine relative to its tetrad plane, we defined an out-of-plane angle by using a pseudo-improper dihedral scheme. The tetrad plane was defined by the C1' atom of the guanine (G) being analyzed and the two C1' atoms of its neighboring guanines (G+ and G-). The O6 atom of G was used to define the out-of-plane angle.
